# All-trans retinoic acid recalibrates macrophage transcriptional responses to drug-resistant *Mycobacterium tuberculosis* through strain-specific immunometabolic reprogramming

**DOI:** 10.64898/2026.04.01.715810

**Authors:** Mona Roshan Singh, Roshan Kumar, Shailly Anand, Isha Pahuja, Md. Arafat Hussain, Roushan Kumar, Archana Singh, Ved Prakash Dwivedi, Manoj Kumar, Akinyemi I. Ojesina, Indrakant Kumar Singh

**Affiliations:** Department of Zoology, Deshbandhu College, University of Delhi, New Delhi, 110019, India; Department of Obstetrics and Gynecology, Division of Gynecologic Oncology, Medical College of Wisconsin, Milwaukee, WI 53226, USA; Post-Graduate Department of Zoology, Magadh University, Bodh Gaya, Bihar, India; Medical College of Wisconsin Cancer Center, Milwaukee, WI, 53226, USA; Department of Zoology, Deen Dayal Upadhyaya College, Sector-3, Dwarka, New Delhi - 110078, India; Immunobiology Group, International Centre for Genetic Engineering and Biotechnology (ICGEB), New Delhi, India; Department of Pulmonary, critical Care and Sleep Medicine, AIIMS, New Delhi-110029, India; Department of Plant Molecular Biology, University of Delhi (South Campus), New Delhi, 110021, India; Environmental Toxicology Group, CSIR-Indian Institute of Toxicology Research (CSIR-IITR), Vishvigyan Bhavan, 31, Mahatma Gandhi Marg, Lucknow 226001, Uttar Pradesh, India; Department of Microbiology and Immunology, Medical College of Wisconsin, Milwaukee, WI, USA; Dr. B. R. Ambedkar Centre for Biomedical Research, University of Delhi, Delhi, 110007, India

**Author notes:** Equal Contribution (Mona Roshan Singh, Roshan Kumar).

## Abstract

Tuberculosis (TB) caused by *Mycobacterium tuberculosis* (*M.tb*) remains the leading infectious cause of death worldwide, with multidrug-resistant (MDR) and extensively drug-resistant (XDR) strains presenting an escalating therapeutic crisis. All-trans retinoic acid (ATRA), an active vitamin A metabolite with established immunomodulatory properties, has emerged as a candidate host-directed therapy, yet its genome-wide transcriptional effects on macrophage across strains of differing drug-resistance profiles remain uncharacterized. We performed RNA sequencing of murine peritoneal macrophages infected with drug-susceptible H37Rv, MDR-2261, or XDR-MCY *M.tb* strains, with and without ATRA treatment, validated by RT-PCR and flow cytometry. All three strains activated a conserved pro-inflammatory program dominated by *Nos2*, *Il1b*, and *Acod1* induction with progressive suppression of host translational and mitochondrial machinery. XDR-MCY uniquely induced *Ifnb1*/IFN-β, suppressed *Prdx1* and *Gpx4*, the latter shared with MDR-2261 and showed selective downregulation of MHC-II processing genes, suggesting a multilayered immune evasion strategy. ATRA activated canonical retinoid signaling across all infection states and consistently induced *Arg1*-mediated resolution signaling, with *Nos2* suppression observed in H37Rv- and MDR-infected macrophages. ATRA selectively restored *Epas1*/HIF-2α in H37Rv- and MDR-infected macrophages without disrupting the HIF-1α-driven antimicrobial program or itaconate biosynthesis. ATRA responsiveness progressively attenuated with drug resistance, with XDR-MCY-infected macrophages largely refractory to transcriptional reprogramming. These findings provide a transcriptional rationale for evaluating ATRA as a host-directed adjunct in drug-resistant TB and identify *Gpx4* suppression in drug-resistant strains and XDR-specific *Ifnb1* induction as vulnerabilities necessitating further investigation.

## Introduction

*Mycobacterium tuberculosis (M.tb)*, the causative agent of TB is a persistent global health threat that has co-evolved with humans, causing significant morbidity and mortality worldwide. According to the World Health Organization (WHO), Global Tuberculosis Report 2025, 10.7 million people developed TB globally in 2024 causing 1.23 million deaths, making it the single leading cause of death from an infectious agent worldwide and one of the top10 cause of death worldwide (1). The emergence of MDR and XDR TB has further exacerbated the challenge of combating this serious global threat, requiring urgent alternative therapeutic strategies (2, 3). Increasing resistance to at least rifampicin and isoniazid, the two most effective first-line drugs used in the traditional treatment regimen of Directly Observed Treatment Short-course (DOTs) for curing TB has not only increased the duration of therapy (4), raised the treatment cost (5), but has also led to severe side effects (6–8) including immunotoxicity, and heightened susceptibility to reinfection. Resistance emergence has been driven by incomplete treatment adherence, delayed diagnosis, inadequate healthcare infrastructure, and overuse or misuse of second-line agents.

ATRA, the active metabolite of Vitamin A, is integral to a variety of essential physiological processes, including growth, development, immune function, and cellular differentiation (9, 10). ATRA plays a significant role in embryonic development and is widely recognized as an effective immunomodulatory agent (11–13). It is particularly noted for its ability to modulate immune responses by enhancing the activity of various immune cells, including T cells, dendritic cells, and macrophages (14–16). This capability has led to growing interest in its therapeutic potential, due to its ability to regulate both innate and adaptive immunity, including the activation of anti-inflammatory pathways and the resolution of inflammation, thereby combating several infections (10, 17).

Over the past several years, there has been an increasing interest in exploring the relationship between ATRA and TB treatment (16–18). This is attributed to the ability of ATRA to modulate immune responses in ways that could strengthen the host defense mechanisms against *M.tb*. Furthermore, deficiency of Vitamin A is associated with 10-fold increased susceptibility of tuberculosis infection (19). Research has demonstrated that ATRA may influence the host’s ability to mount an effective immune response, which could contribute to the control of infection and potentially shorten the duration of therapy (16). Moreover, its ability to impact granuloma formation, a key feature in the immune response to TB, has added another layer of intrigue to its potential role in TB treatment (20). Despite this promise, the genome-wide transcriptional mechanisms by which ATRA recalibrates macrophage responses to drug-resistant *Mtb* strains remain poorely understood, and it is still unclear whether its immunomodulatory effects vary across drug-susceptible, MDR, and XDR strains.

Through this transcriptomic study, we investigate the immunomodulatory effects of ATRA on macrophage responses to *M. tb* infection. We examined macrophage responses to three *M. tb* strains (RV, MDR, and XDR) with and without ATRA treatment, analyzing transcriptional changes 24 hours post-treatment. This study leverages comprehensive pathway analysis to characterize strain specific host responses and ATRA mediated immune modulation, thereby providing a mechanistic framework to evaluate ATRA as a potential host-directed adjunct in drug-resistant TB.

## Results

### Experimental Setup and RNA-seq Data Generation

The experiment was performed to investigate the transcriptional effects of ATRA treatment on macrophages infected with different *M.tb* strains (H37Rv, MDR-2261, and XDR-MCY). Thioglycolate-induced peritoneal macrophages from C57BL/6 mice were isolated and infected with distinct *M.tb* strains (Fig. 1A). Eight experimental groups were established: uninfected controls (UNINF), uninfected with ATRA treatment (UNINF_RA), H37RV strain infection (RV), H37RV strain infection with ATRA (RVATRA), multidrug-resistant strain infection (MDR-2261), MDR with ATRA treatment (MDR-RA), extensively drug-resistant strain infection (XDR-MCY), and XDR with ATRA treatment (XDR-RA). Each group comprised of three technical replicates. RNA sequencing was thus performed on 24 samples of which one sample from the UNINF_RA group was excluded due to poor quality. Overall, the sequencing depth ranged from 43.5 to 67.5 million raw reads/sample, yielding 6.5 to 10.1 GB of raw data (Supplementary Table 1). High-quality sequencing was achieved across all samples (except UNINF_RA), with effective rates of 94.9–98.98% and Q30 scores of 92.43–95.08%. The PCA and Poisson distance heatmap reveal that infection status is the dominant driver of transcriptional variance (64%), with samples clustering distinctly by *M.tb* strain (Fig. 1B). While ATRA treatment consistently modulates expression across all conditions, its secondary impact (11% variance) is most pronounced in XDR-infected and uninfected macrophages (Fig. 1 B & C).

**Figure 1:**
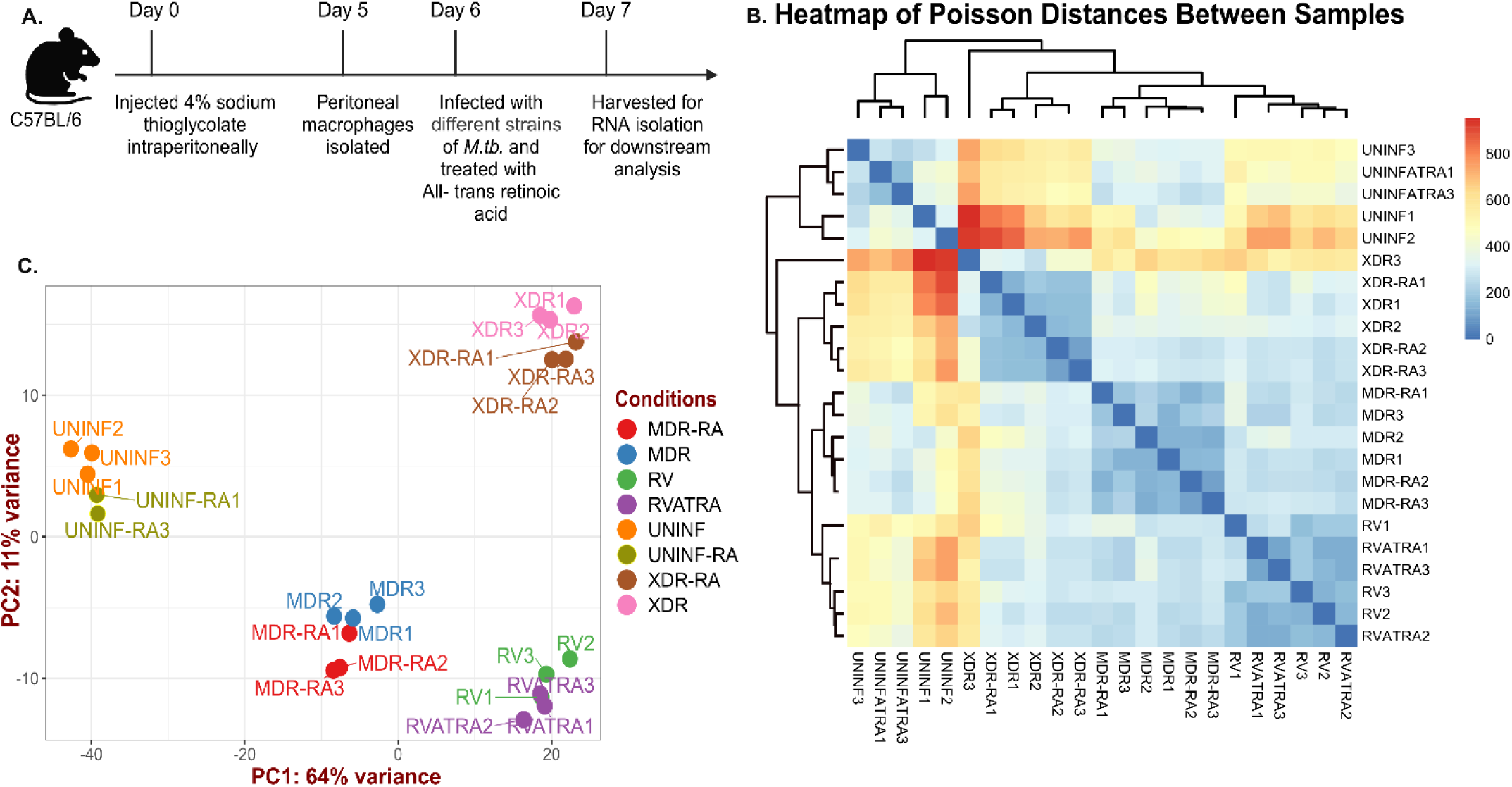
Transcriptional Response of Macrophages to *M. tb* Infection and Retinoic Acid Treatment. **A. Experimental Design:** C57BL/6 mice were injected intraperitoneally with 4% sodium thioglycolate to induce peritonitis. Peritoneal macrophages were isolated on day 5, infected with different strains of *M.tb* (H37RV, MDR-2261, XDR-MCY), and treated with all-trans retinoic acid (RA) or left untreated. Macrophages were harvested on day 7 for RNA isolation and downstream analysis. **B.** Poisson distance correlation heatmap showing the pairwise correlation between samples based on Poisson distance. Samples are clustered using complete linkage **C.** Principal Component Analysis (PCA) plot showing the clustering of samples based on gene expression profiles. Samples are color-coded by condition. PC1 and PC2 explain 64% and 11% of the variance, respectively.

### Differential Gene Expression Changes in Response to Infection

The differential gene expression analysis revealed extensive and strain-dependent gene expression reprogramming in response to *M. tb* infection. A total of 2,846 unique differentially expressed genes (DEGs; L2FC I1.5I, FDR<0.05) were identified across all three pairwise comparisons (each strain vs uninfected), of which 1,291 were upregulated and 1,555 downregulated (Fig. 2A & B; Supplementary Table 2).

**Figure 2.**
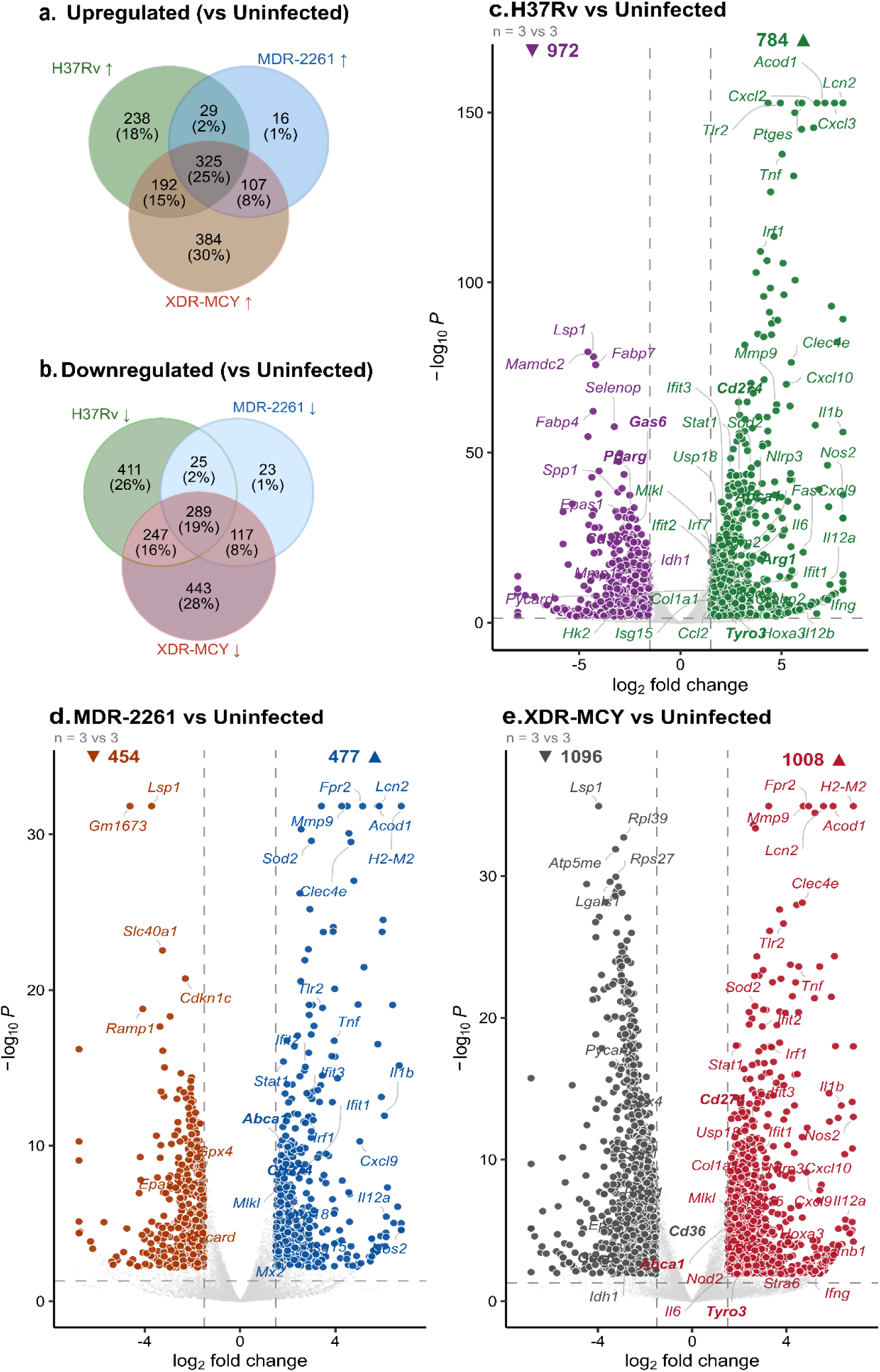
Transcriptional reprogramming of macrophages in response to M. tb infection across drug-resistance strains. (a, b) Venn diagrams showing the overlap of significantly upregulated (a) and downregulated (b) genes among macrophages infected with H37Rv (green), MDR-2261 (blue), and XDR-MCY (red) compared to uninfected controls. A core set of 325 upregulated and 289 downregulated genes was commonly regulated across all three strains, representing 25% and 19% of the total union, respectively. (c–e) Volcano plots depicting differential gene expression in H37Rv-- (c), MDR-2261-- (d), and XDR-MCY-- infected (e) macrophages versus uninfected controls (n = 3 per group). Numbers in the upper corners indicate the total DEG counts for each direction. Dashed vertical lines indicate the |log₂FC| ≥ 1.5 threshold while; the dashed horizontal line indicates padj = 0.05.

In H37Rv-infected macrophages (RV vs UNINF), 1,756 DEGs were identified (784 upregulated, 972 downregulated) (Fig. 2C; Supplementary Table 2). The most highly upregulated genes included *Nos2* (L2FC= 14.03), *Lcn2* (L2FC= 8.25), *Cxcl3* (L2FC= 7.61), *Acod1* (L2FC= 7.14), and *Cxcl2* (L2FC= 6.74), reflecting strong activation of antimicrobial and pro-inflammatory programs. Key immune mediators were also substantially induced, including *Il1b* (L2FC= 10.17), *Il12a* (L2FC= 7.39), *Il12b* (L2FC= 6.16), *Tnf* (L2FC= 5.02), *Il6* (L2FC= 3.41), *Cxcl9* (L2FC= 6.07), and *Cxcl10* (L2FC= 5.23). Among the most significantly downregulated genes were *Mamdc2* (L2FC= −4.57), *Lsp1* (L2FC= −4.30), *Fabp4* (L2FC= −4.31), *Fabp7* (L2FC= −4.19), and *Selenop* (L2FC= −3.26). of note, downregulation was also observed for lipid metabolism regulators *Pparg* (L2FC= −2.43) and *Cd36* (L2FC= −2.62), along with the ECM-associated gene *Spp1* (L2FC= −3.09).

In MDR-2261-infected macrophages (MDR vs UNINF), 931 DEGs were identified (477 upregulated, 454 downregulated) (Fig. 2D; Supplementary Table 2). The top upregulated genes included H2-*M2* (L2FC= 7.43), *Lcn2* (L2FC= 5.85), *Acod1* (L2FC= 5.79), *Fpr2* (L2FC= 5.14), and *Mmp9* (L2FC= 4.49). Downregulated genes included *Gm1673* (L2FC= −4.62), *Ramp1* (L2FC= −4.09), *Lsp1* (L2FC= −3.71), *Slc40a1* (L2FC= −3.25), and *Cdkn1c* (L2FC= −2.30).

XDR-MCY infection elicited the broadest transcriptional response, with 2,104 DEGs identified (1,008 upregulated, 1,096 downregulated) (Fig. 2D; Supplementary Table 2). The most highly upregulated genes included *H2-M2* (L2FC= 7.73), *Acod1* (L2FC= 5.98), *Lcn2* (L2FC= 5.57), *Fpr2* (L2FC= 4.94), and *Mmp9* (L2FC= 4.71). The top downregulated genes were *Lsp1* (L2FC= −3.97), *Lgals1* (L2FC= −3.49), *Atp5me* (L2FC= −3.25), *Rps27* (L2FC= −3.24), and *Rpl39* (L2FC= −2.90), suggesting suppression of cellular translation and oxidative metabolism. Across all three comparisons, *Acod1*, *Lcn2*, and *Lsp1* consistently emerged among the top dysregulated genes.

### Differential Pathways in Response to M. tb strains Infection

To systematically characterize the biological processes activated or suppressed in macrophages following infection with different *M. tb* strains, IPA canonical pathway analysis was performed on DEGs from each strain-specific comparison (H37Rv, MDR-2261, and XDR-MCY, each versus uninfected controls) and on the intersecting gene set shared across all three infected conditions (Fig. 3). A total of 123, 156, 145, and 179 significantly dysregulated pathways were identified in the core, H37Rv, MDR-2261, and XDR-MCY comparisons respectively (|z-score| ≥ 2, p ≤ 0.05)(Fig. 3A-D), revealing both a conserved host response to mycobacterial infection and strain-specific transcriptional signatures that escalate in magnitude with increasing drug resistance.

**Figure 3.**
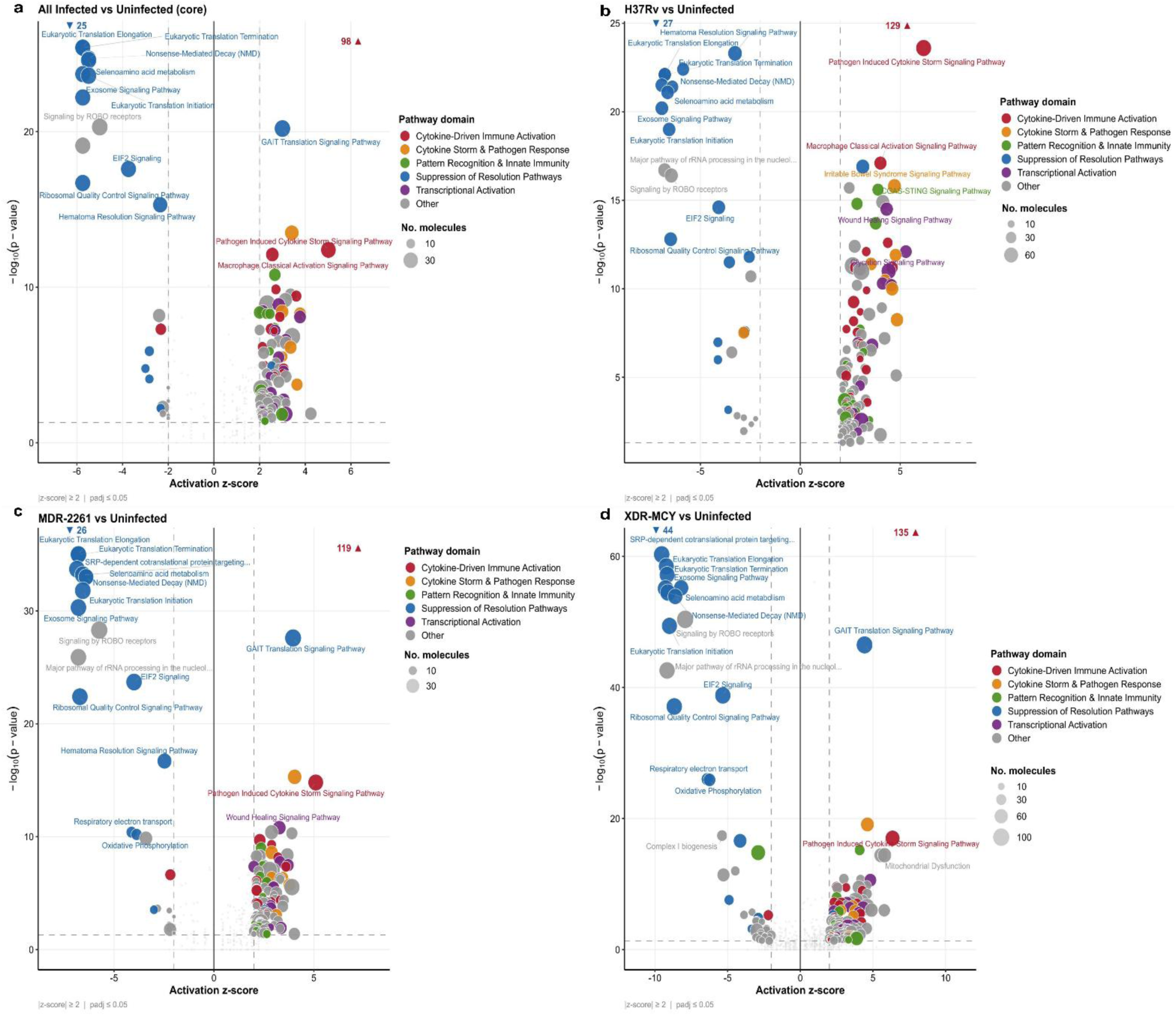
IPA canonical pathway analysis of macrophage transcriptional responses to M. tb infection. (a) Pathway analysis of core differentially expressed genes shared across all three infected strains versus uninfected, (b–d) Strain-specific pathway analysis comparing H37Rv- (b), MDR-2261- (c), and XDR-MCY-infected (d) macrophages against uninfected controls (n = 3 per group). Each dot represents one IPA canonical pathway. Position on the x-axis indicates the activation z-score, where positive values denote pathway activation and negative values denote inhibition relative to the uninfected condition. The y-axis shows the −log₁₀(p-value) of pathway enrichment. Dot size is proportional to the number of dataset molecules mapping to the pathway. Dot colour indicates the biological domain: red, cytokine-driven immune activation; orange, cytokine storm and pathogen response; green, pattern recognition and innate immunity; blue, suppression of resolution pathways; purple, transcriptional activation; grey, other. Dashed vertical lines indicate |z-score| = 2; the dashed horizontal line indicates p = 0.05. Pathway analysis was performed using Ingenuity Pathway Analysis (IPA, QIAGEN).

### Core Pathways

Analysis of the 614 DEGs constituting the core gene set shared across all three infected conditions identified 98 activated and 25 inhibited pathways, providing a comprehensive macrophage response to *M.tb* irrespective of drug resistance (Fig. 3A; Supplementary Table 3). Among the activated pathways, the *Pathogen Induced Cytokine Storm Signaling Pathway* was the most prominently enriched (z-score = 5.01), driven by coordinated upregulation of a broad network of pro-inflammatory mediators (*Ccl3, CCL5, CXCL9, CXCL13, IL1A, IL1B, IL12A, TNF, CSF2, NOS2*). This was supported by robust activation of *Macrophage Classical Activation Signaling* (z-score = 2.56) and the *CGAS-STING Signaling Pathway* (z-score = 2.67), indicating that cytosolic DNA sensing and type I interferon induction represent core, strain-independent features of the macrophage *M. tb* interaction. Innate immune signaling pathways were broadly engaged, including *Toll-like Receptor Signaling* (z-score = 2.45), *NOD1/2 Signaling Pathway* (z-score = 2.00), and *Pattern Recognition Receptor Signaling* (z-score = 2.65), confirming comprehensive activation of both surface and intracellular pathogen detection mechanisms. Interestingly, the *GAIT (Gamma-Interferon–Activated Inhibitor of Translation) Translation Signaling Pathway* was highly activated (z-score = 3.00) in the core gene set, indicating selective translational control of interferon-γ-activated inhibitor of translation as part of the post-transcriptional regulatory response. *Interleukin-10 signaling* (z-score = 2.716) and *Interferon α/β Signaling* (z-score = 2.31) were also activated in the shared response, suggesting simultaneous engagement of both pro-inflammatory amplification and counter-regulatory type I interferon cascades.

In contrast to this inflammatory activation, the inhibited pathway signature was defined by near-complete suppression of cellular translational machinery, representing the most statistically robust signal across all comparisons (Fig. 3A; Supplementary Table 3). *Eukaryotic Translation Elongation and Termination* (both z-score = −5.75, −log₁₀p = 25.4), *Nonsense-Mediated Decay* (z-score = −5.48, −log₁₀p = 24.7), *Response of EIF2AK4 to Amino Acid Deficiency* (z-score = −5.48, −log₁₀p = 24.7), *Selenoamino Acid Metabolism* (z-score = −5.49, −log₁₀p = 24.6), *SRP-Dependent Co-translational Protein Targeting to Membrane* (z-score = −5.75, −log₁₀p = 23.7), *Eukaryotic Translation Initiation* (z-score = −5.49, −log₁₀p = 23.6), and *Exosome Signaling* (z-score = −5.75, −log₁₀p = 22.2) were all inhibited, collectively suggesting a comprehensive shutdown of the ribosomal and translational apparatus. *Major Pathway of rRNA Processing* (z-score = −5.75), Ribo*somal Quality Control* (z-score = −5.75), and *EIF2 Signaling* (z-score = −3.74) further emphasized this suppression of host cellular translational machinery. The oxidative Phosphorylation (z-score = −2.83) was also inhibited in the core gene set, indicating that suppression of mitochondrial energy metabolism is an additional feature of macrophage infection by *M. tb*.

### H37Rv-Specific Pathway

Comparison of H37Rv-infected macrophages with uninfected controls identified 128 activated and 27 inhibited pathways (Fig. 3B; Supplementary Table 4). The *Pathogen Induced Cytokine Storm Signaling Pathw*ay reached its highest enrichment in this comparison (z-score = 6.17), reflecting the large number of cytokine and chemokine genes upregulated by drug-susceptible H37Rv. *Macrophage Classical Activation Signaling* (z-score = 4.01) was the second most enriched activation pathway, confirming a canonical M1 polarization response. The *CGAS-STING Signaling Pa*thway was strongly activated (z-score = 3.89), consistent with robust cytosolic nucleic acid sensing during productive H37Rv infection, H37Rv infection specifically activated the *Glycation Signaling Pathway* (z-score = 5.28) and the *Cachexia Signaling Pathway* (z-score = 4.84), pointing to metabolic reprogramming and tissue catabolism pathways that may contribute to the systemic wasting phenotype observed in tuberculosis. *IL-17A Signaling in Fibroblasts* (z-score = 4.38), *NOD1/2 Signaling* (z-score = 3.77) were also among the top activated pathways, highlighting the innate immune receptor engagement and downstream inflammatory amplification.

The inhibited pathway profile in H37Rv infection was characterized by the most extreme z-scores among all comparisons, with *Eukaryotic Translation Elongation and Termi*nation (z-score = −6.78), *SRP-Dependent Co-translational Protein Targeting* (z-score = −6.93), *Nonsense-Mediated Decay* (z-score = −6.40), and *Selenoamino Acid Metabolism* (z-score = −6.64) all exhibiting near-maximal inhibition (Fig. 3B; Supplementary Table 4). The *Hematoma Resolution Signaling Pathway* was significantly inhibited (z-score = −3.27, −log₁₀p = 23.3), and was particularly the most statistically significant inhibited pathway in the H37Rv comparison, suggesting that suppression of tissue repair programs is particularly pronounced during drug-susceptible infection. H37Rv infection additionally inhibited the *Tuberculosis Active Signaling Pathway* (z-score = −2.83), *FXR/RXR Activation* (z-score = −3.55), and *Erythropoietin Signaling Pathway* (z-score = −2.5). The suppression of FXR/RXR signaling pathway suggests lipid metabolic disruption and nuclear receptor-mediated anti-inflammatory responses being inhibited during infection.

### MDR-2261-Specific Pathway

In MDR-2261-infected macrophages, 119 activated and 26 inhibited pathways were identified (Fig. 3c, Supplementary Table 5). Interestingly, the *GAIT Translation Signaling Pathway* emerged as the most statistically significant activated pathway in MDR-2261 infection (z-score = 3.96, −log₁₀p = 27.6), surpassing the *Pathogen Induced Cytokine Storm pathway* (z-score = 5.10, −log₁₀p = 14.8) in terms of significance. This finding suggests that selective translational reprogramming of cytokine-related transcripts via the GAIT complex is a prominent feature of the macrophage response to drug-resistant MDR-2261, potentially representing an adaptive counter-regulatory mechanism. The *Wound Healing Signaling Pathway* (z-score = 3.27) indicates simultaneous tissue remodeling responses alongside inflammatory activation. CGAS-STING Signaling was upregulated (z-score = 2.36), suggesting attenuated cytosolic nucleic acid sensing in MDR-2261-infected macrophages.

The inhibited pathway profile in MDR-2261 infection was more severe than in H37Rv, with all translational pathway z-scores reaching the same extreme values (*Eukaryotic Translation Elongation/Termination* z-score = −6.78; *SRP-Dependent Co-translational Protein Targeting* z-score = −6.86) (Fig. 3c, Supplementary Table 5), but at substantially higher significance, indicating greater statistical depth of translational suppression with a larger proportion of the ribosomal proteome affected. Interestingly, MDR-2261 infection additionally inhibited *Respiratory Electron Transport* (z-score = −4.12, −log₁₀p = 10.4) and Oxidative Phosphorylation (z-score = −3.87, −log₁₀p = 10.2), pathways, indicating that drug-resistant infection extends macrophage metabolic suppression from the ribosome to the mitochondrial electron transport chain. Furthermore, *Complex I Biogenesis* (z-score = −2.83, −log₁₀p = 3.64) and *Complex III Assembly* (z-score = −2.24) were also downregulated, confirming abruption in complete oxidative phosphorylation assembly pathways. This metabolic burden imposed by MDR-2261 may reflect either enhanced bacterial strategies to subvert macrophage bioenergetics or a more severe host immune metabolic response.

### XDR-MCY-Specific Pathway

XDR-MCY infection resulted in the most extensive and pronounced pathway dysregulation among all comparisons, with 135 activated and 44 inhibited (Fig. 3D; Supplementary Table 6). The depth and magnitude of inhibition were qualitatively distinct from other comparisons. The *GAIT Translation Signaling Pathway* again ranked as the most statistically significant activated pathway (z-score = 4.42, −log₁₀p = 46.5). The *Pathogen Induced Cytokine Storm Signaling Pathway* showed the highest activation z-score across all comparisons (z-score = 6.35), consistent with XDR-MCY eliciting the most severe inflammatory cascade. The *Mitochondrial Dysfunction Pathway* was also activated (z-score = 5.80), likely reflecting compensatory mitochondrial stress signaling and reactive oxygen species-driven gene expression in response to severe energetic disruption. XDR-MCY infection also activated pathways related to extracellular matrix remodeling and structural tissue changes: *Assembly of Collagen Fibrils and Multimeric Structures* (z-score = 4.15, −log₁₀p = 10.7), *Collagen Biosynthesis and Modifying En*zymes (z-score = 4.58, −log₁₀p = 9.78), *Extracellular Matrix Organization* (z-score = 4.60, −log₁₀p = 8.50), and *Integrin Cell Surface Interactions* (z-score = 3.00, −log₁₀p = 10.84), suggesting that XDR-MCY infection drives pathological tissue remodeling and fibrotic responses in macrophages. *Granzyme A Signaling* (z-score = 4.08, −log₁₀p = 15.2) was specifically activated in XDR infection, indicative of cytotoxic effector pathway engagement, while Interleukin-4 and Interleukin-13 Signaling (z-score = 2.5) suggested activation of Th2-associated counter-regulatory pathway not observed in other strains.

The inhibited pathway profile in XDR-MCY infection was categorically more extreme than in any other comparison (Fig. 3D; Supplementary Table 6). *SRP-Dependent Co-translational Protein Targeting* (z-score = −9.54, −log₁₀p = 60.3), *Eukaryotic Translation Elongation* (z-score = −9.22, −log₁₀p = 58.5), *Eukaryotic Translation Termination* (z-score = −9.16, −log₁₀p = 57.2), *Response of EIF2AK4 to Amino Acid Deficiency* (z-score = −8.20, −log₁₀p = 55.2), and *Exosome Signaling* (z-score = −9.27, −log₁₀p = 55.1) were inhibited at levels that substantially exceed those seen in MDR-2261 and H37Rv infections. Mitochondrial pathway inhibition was also comprehensive in XDR infection: *Respiratory Electron Transport* (z-score = −6.40, −log₁₀p = 26.0), *Oxidative Phosphorylation* (z-score = −6.25, −log₁₀p = 25.9), Co*mplex I Biogenesis* (z-score = −5.38, −log₁₀p = 17.4), *Complex IV Assembly* (z-score = −4.47, −log₁₀p = 12.0), and *Mitochondrial Translation* (z-score = −4.90, −log₁₀p = 7.57), representing a complete shutdown of the mitochondrial respiratory chain assembly and function. The *Neutrophil Extracellular Trap Signaling Pathway* (z-score = −2.90) was uniquely inhibited in XDR infection, suggesting active suppression of NET formation mechanisms that may contribute to immune evasion by this extensively drug-resistant strain. The *Stress Granule Signaling Pathway* reached its highest inhibition level in XDR infection (z-score = −5.29, −log₁₀p = 11.4), consistent with acute disruption of cellular stress response mechanisms.

### Comparative Pathway Landscape Across Strains

The comparative pathway analysis reveals a hierarchical and quantitatively escalating pattern of macrophage pathway dysregulation that correlates with increasing drug resistance. All three strains converge on activation of cytokine storm, classical macrophage activation, GAIT translation control, and innate pattern recognition pathways, while universally suppressing the translational machinery and resolution programs. However, the magnitude of both activation (z-scores escalating from H37Rv to XDR for cytokine storm) and inhibition (ribosomal pathway z-scores increasing from approximately −6.8 in H37Rv to −9.5 in XDR) increases progressively. Infections with MDR-2261 and XDR-MCY additionally induce mitochondrial bioenergetic suppression that is not pronounced in H37Rv infection, while XDR-MCY activated extracellular matrix remodeling and cytotoxic signaling pathways more comprehensively and uniquely inhibited NET formation. These strain-specific differences suggest that acquired drug resistance in *M.tb* is accompanied by enhanced capacity to broadly disrupt macrophage metabolic homeostasis, extending the host transcriptional impact from translational suppression alone (H37Rv) to encompass mitochondrial energy production, structural tissue remodeling, and cytotoxic effector mechanisms (XDR-MCY). The universal inhibition of Hematoma Resolution Signaling across all comparisons further indicates that *M. tb* infection systematically impairs macrophage resolution capacity irrespective of strain, potentially prolonging the inflammatory microenvironment required for bacterial persistence.

#### Differential Gene Expression in Response to ATRA Treatment Post-Infection

To identify the common transcriptional effects of ATRA common across all infection states, a strain-corrected pooled analysis was performed comparing ATRA-treated infected macrophages (RVATRA + MDR-RA + XDR-RA; n = 9) against untreated infected macrophages (RV + MDR + XDR; n = 9), with bacterial strain included to prevent confounding. This analysis identified 44 DEGs: 31 upregulated and 13 downregulated (Supplementary Table 2). The most consistently and strongly upregulated genes were the retinoic acid-metabolizing enzymes *Cyp26b1* (L2FC= 5.42) and *Cyp26a1* (L2FC= 3.19), alongside the canonical RAR target *Rarb* (L2FC= 2.04) and the retinol transporter *Stra6* (L2FC= 2.06), confirming robust activation of the retinoic acid receptor signaling axis across all infection backgrounds. *Mfng* (L2FC= 2.33) and *Medag* (L2FC= 1.87) were also consistently upregulated. Among downregulated genes, the BMP antagonist *Grem1* showed the greatest decrease (L2FC= −2.38), followed by the prostaglandin-catabolizing enzyme *Hpgd* (L2FC= −2.06), *Sfrp2* (L2FC= −2.10), *Cck* (L2FC= −1.78), and *Igfbp2* (L2FC= −1.68).

Analysis of per-strain ATRA responses revealed different magnitudes of transcriptional reprogramming depending on the infecting strain (Fig. 4D-F). In H37Rv-infected macrophages, ATRA treatment resulted in 103 DEGs (51 upregulated, 52 downregulated). Upregulated genes included *Mfng* (L2FC= 2.31), *Epas1* (L2FC= 1.91), and *Cxcl3* (L2FC= 1.89), while *Nos2* (L2FC= −1.63) and *Cxcl10* (L2FC= −1.93) were significantly downregulated (Fig. 4D), suggesting that ATRA attenuates select pro-inflammatory effector programs even in classically activated macrophages. In MDR-2261-infected macrophages, 68 DEGs were identified (49 upregulated, 19 downregulated), with the strongest induction again in the retinoid pathway: *Cyp26b1* (L2FC= 5.66), *Cyp26a1* (L2FC= 3.48), and *Camkk1* (L2FC= 3.77) (Fig. 4E). of note, *Arg1* was strongly upregulated by ATRA specifically in the MDR infection context (L2FC= 4.13), suggesting that ATRA promotes a shift toward alternative macrophage activation in drug-resistant infection. In contrast, ATRA treatment of XDR-MCY-infected macrophages yielded only 9 DEGs (7 upregulated, 2 downregulated) (Fig. 4F). Upregulated genes were limited to retinoid pathway (*Cyp26b1*, L2FC= 4.51; *Cyp26a1*, L2FC= 3.32) and pseudogenes, with *Sfrp2* (L2FC= −1.98) and *Grin2a* (L2FC= −5.04) as the only downregulated genes. This near-absent transcriptional response suggests that macrophages are substantially less permissive to ATRA-mediated transcriptional reprogramming during XDR-MCY infection compared to H37Rv or MDR-2261 infection.

**Figure 4.**
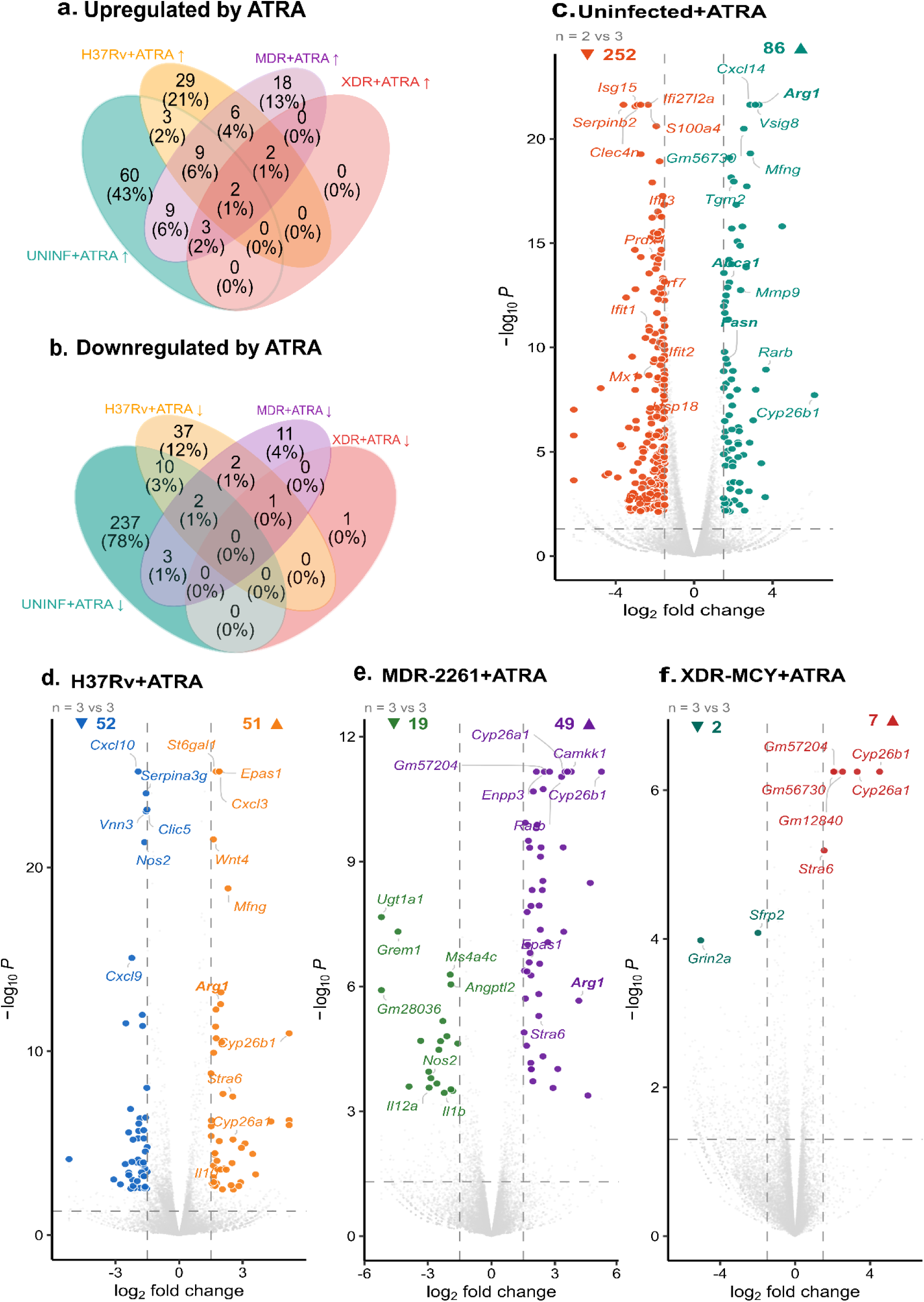
Transcriptional effects of ATRA treatment across *M. tb* infection states. (a, b) Venn diagrams showing the overlap of significantly upregulated (a) and downregulated (b) genes in response to ATRA treatment across four conditions: Uuninfected (UNINF+ATRA), H37Rv-infected (H37Rv+ATRA), MDR-2261-infected (MDR+ATRA), and XDR-MCY-infected (XDR+ATRA) macrophages, each compared to their respective untreated counterparts. Numbers indicate DEG counts with percentages of the total union shown in parentheses. (c) Volcano plot depicting differential gene expressions in uninfected macrophages treated with ATRA versus untreated uninfected macrophages (n = 2 vs 3). (d–f) Volcano plots depicting the effect of ATRA treatment effect in H37Rv- (d), MDR-2261- (e), and XDR-MCY-infected (f) macrophages versus their respective untreated infected counterparts (n = 3 vs 3 for all). DEG counts are indicated in the upper corners. Dashed vertical lines indicate the |log₂FC| ≥ 1.5 threshold; the dashed horizontal line indicates padj = 0.05.

In uninfected macrophages, ATRA treatment produced the broadest response overall, yielding 338 DEGs (86 upregulated, 252 downregulated; n = 2 vs 3) (Fig. 4C). The predominance of downregulated genes (252 vs 86 upregulated) contrasted with the primarily upregulatory ATRA effect seen in infected contexts and may reflect suppression of basal inflammatory tone in the absence of pathogen-derived signals. Among upregulated genes, *Arg1* (L2FC= 3.30), *Vsig8* (L2FC= 3.13), *Mfng* (L2FC= 2.87), *Cxcl14* (L2FC= 2.85), and the retinoid targets *Cyp26b1* (L2FC= 6.60) and *Rarb* (L2FC= 3.66) were most significantly induced. The most prominently downregulated genes were *Serpinb2* (L2FC= −3.60), *Isg15* (L2FC= −2.88), *Clec4n* (L2FC= −2.85), *Ifi27l2a* (L2FC= −2.73), and *S100a4* (L2FC= −2.35). Notably, multiple interferon-stimulated genes were significantly suppressed, including *Ifit1* (L2FC= −2.30), *Ifit2* (L2FC= −1.99), *Ifit3* (L2FC= −2.14), *Mx1* (L2FC= −2.06), and *Usp18* (L2FC= −1.54), indicating that ATRA broadly attenuates interferon-driven gene expression even in the absence of active infection.

Further, analysis of the ATRA response across all four conditions (Fig. 3a, b) confirmed that the transcriptional effects of ATRA are largely context-specific. Of the total 141 upregulated genes across all ATRA comparisons, 60 (43%) were unique to the uninfected condition, 29 (21%) to H37Rv-infected, and 18 (13%) to MDR-infected macrophages, with no genes uniquely upregulated in the XDR-MCY context. Only 2 genes were commonly upregulated across all four ATRA comparisons. A similar pattern was observed for downregulated genes: of a total 304, 237 (78%) were unique to uninfected ATRA-treated macrophages, with no genes commonly downregulated across all four conditions. Together, these findings demonstrate that while ATRA consistently activates its canonical retinoid receptor target gene program across infection states, the broader transcriptional impact of ATRA is highly dependent on the underlying macrophage activation state imposed by the infecting *M. tb* strain.

### Pathway Analysis in Response to Retinoic Acid Treatment Post-Infection

To characterize the biological pathways underlying the transcriptional reprogramming induced by ATRA treatment, we performed IPA canonical pathway analysis on DEGs from each ATRA versus untreated comparison across four macrophage backgrounds: uninfected (UNINF+ATRA vs UNINF), H37Rv-infected (H37Rv+ATRA vs H37Rv), MDR-2261-infected (MDR+ATRA vs MDR), and XDR-MCY-infected (XDR+ATRA vs XDR) macrophages.

### ATRA Effect in Uninfected Macrophages

In the absence of infection, ATRA treatment elicited the most extensive pathway-level response, with 23 pathways showing significant dysregulation, including 4 activated and 19 inhibited (Supplementary Fig. 1 & Supplementary Table 7). This pronounced dominance of pathway inhibition over activation was a distinctive feature of the uninfected ATRA response, in contrast to the primarily activating effect of ATRA observed in infected contexts. The activated pathways were modest in number but biologically meaningful. RAR Activation was significantly enriched (z-score=2.83), confirming on-target engagement of the canonical retinoic acid receptor signaling axis and validating the biological activity of ATRA in this experimental system.

The inhibited pathway profile revealed suppression of ribosomal and translational machinery, mirroring the pattern observed during infection but in this instance driven by ATRA rather than a bacterial challenge. *SRP-Dependent Co-translational Protein Targeting to Membrane* (z-score = −3.87), *Eukaryotic Translation Elongation and Termination* (both z-score = −3.74), *Selenoamino Acid M*etabolism (z-score = −3.74), *Nonsense-Mediated Decay* (z-score = −3.46), *Response of EIF2AK4 to Amino Acid Deficiency* (z-score = −3.46), *Eukaryotic Translation Initiation* (z-score = −3.74), and *Exosome Signaling Pathway* (z-score = −3.74) were all significantly inhibited, indicating that ATRA recapitulates translational suppression in uninfected macrophages in a manner mechanistically overlapping with the infection response. *EIF2 Signaling* (z-score = −2.83, −log₁₀p = 7.88), *Signaling by ROBO Receptors* (z-score = −3.87), *Major Pathway of rRNA Processing* (z-score = −3.74), and *Ribosomal Quality Control Signaling Pathway* (z-score = −3.74) reinforced the depth of translational suppression. Beyond translational pathways, ATRA inhibited Interferon α/β Signaling (z-score = −2.65), indicating active suppression of the type I interferon response in the absence of infection, a finding consistent with known immunomodulatory properties of retinoic acid that contain basal interferon levels. Further inhibited pathways included *Cytoprotection by HMOX1*, *Oxidative Phosphorylation*, *Respiratory Electron Transport*, *Mitochondrial Protein Import*, *TP53 Regulates Metabolic Genes*, and *ISGylation Signaling Pathway*, collectively indicating that ATRA in uninfected macrophages broadly dampens antioxidant defense, mitochondrial biosynthesis, metabolic stress sensing, and innate immune interferon-related ISGylation processes. The suppression of ISGylation and type I interferon signalling is particularly notable as it indicates that ATRA tempers baseline innate immune activation states in macrophages even prior to pathogenic challenge.

### ATRA Effect in H37Rv-Infected Macrophages

In contrast to the extensive pathway remodeling observed in uninfected macrophages, ATRA treatment of H37Rv-infected macrophages identified only a single significant pathway: *Macrophage Alternative Activation Signaling Pathway* (z-score = 2.00, −log₁₀p = 2.80) involving genes namely *ARG1, EPAS1, IL10, and NOS2*. No inhibited pathways met the threshold of statistical significance. The selective activation of the *Macrophage Alternative Activation Signaling Pathway* indicates that ATRA promotes a shift towards an M2-like macrophage phenotype in the context of H37Rv infection, characterized by anti-inflammatory cytokine production, enhanced efferocytosis capacity, and immunomodulatory gene expression. This is consistent with the known capacity of retinoic acid to promote regulatory and tolerogenic macrophage programs and may represent a mechanism by which ATRA modulates the excessive pro-inflammatory response to drug-susceptible *M.tb* without globally suppressing antimicrobial effector functions.

### ATRA Effect in MDR-2261-Infected Macrophages

ATRA treatment of MDR-2261-infected macrophages identified three significant pathways of which two were activated and one was inhibited. *Macrophage Alternative Activation Signaling Pathway* was the most significantly enriched (z-score = 2.24), recapitulating the H37Rv+ATRA finding and suggesting that ATRA-driven M2-like polarization is a consistent feature of the retinoic acid response in infected macrophages regardless of drug-resistance profile. *RAR Activation* was also identified as a significantly enriched activated pathway (z-score = 2.45), confirming on-target engagement pf retinoid receptors in the MDR infection, thereby validating that the canonical RAR transcriptional response remain intact despite the broader transcriptional dominance of drug-resistant infection.

### ATRA Effect in XDR-MCY-Infected Macrophages

No significant canonical pathways were identified in the XDR+ATRA versus XDR comparison, with zero pathways meeting the threshold (|z-score| ≥ 2.0 and −log₁₀p ≥ 1.3). The suppression of translational machinery, mitochondrial bioenergetics, and stress response pathways by XDR-MCY infection may create a transcriptional environment in which the relatively modest ATRA induced transcriptional response is insufficient to drive enrichment of canonical pathways above the significance threshold.

### ATRA Induces a Comprehensive Efferocytosis Program in Uninfected Macrophages

While apoptosis of infected macrophages has been recognized as a host defense mechanism that limits intracellular mycobacterial replication, apoptosis itself is not sufficient to kill *M.tb*. Bacterial killing requires the subsequent efferocytotic clearance of apoptotic bodies by neighboring phagocytes. Studies have shown that efferocytosis is required to mediate control of the bacterium following apoptosis. The efferocytotic phagosome containing *M. tb* readily fuses with lysosomes, and this leads to bacterial killing both *in vitro* and *in vivo* (21). This positions efferocytosis as the terminal bactericidal checkpoint in a two-step host defence programme: apoptosis limits bacterial replication, but it is efferocytosis that delivers the bacteria to the lysosome for destruction. Understanding whether *M. tb* infection actively suppresses macrophage efferocytosis capacity, and whether this suppression can be pharmacologically reversed, is therefore of direct therapeutic relevance.

To characterize the transcriptional state of the efferocytosis program during active *M. tb* infection and to determine whether ATRA rescues this program, 23 efferocytosis-associated genes spanning all functional modules of the cascade were profiled: TAM receptors (*Mertk*, *Axl*, *Tyro3*), bridging molecules (*Gas6*, *Mfge8*, *Stab2*), engulfment complex (*Elmo1*, *Elmo2*, *Dock1*, *Rac1*), autophagy-lysosomal machinery (*Becn1*, *Atg7*, *Lamp1*), lipid handling (*Abca1*, *Abcg1*, *Cd36*, *Fasn*), nuclear receptors (*Nr1h3*, *Pparg*), resolution mediators (*Arg1*, *Tgfb1*, *Trem2*), and immune checkpoint (*Cd274*), across all seven experimental comparisons by DESeq2 RNA-seq analysis (L2FC| ≥ 1.5, padj < 0.05).

Infection with H37Rv produced a distinctly mixed efferocytosis signature. Four genes were significantly upregulated: *Cd274*/PD-L1 (L2FC= 2.90, 7.5-fold), *Arg1* (L2FC= 2.58, 6.0-fold), *Abca1* (L2FC= +2.50, 5.7-fold), and *Tyro3* (L2FC= +2.44, 5.4-fold). In contrast, three canonical efferocytosis genes were significantly suppressed: *Cd36* (L2FC= −2.62, −6.2-fold), *Gas6* (L2FC= −2.13, −4.4-fold), and *Pparg* (L2FC= −2.43, −5.4-fold). The induction of *Cd274* and immune checkpoint signaling alongside suppression of *Cd36*, the primary macrophage receptor for phosphatidylserine-exposing apoptotic cells, reflects a pattern consistent with active bacterial subversion: *M. tb* appears to upregulate components that dampen immune activation while dismantling the receptor machinery required to recognize and engulf apoptotic debris.

MDR-2261 infection produced a more restricted disruption, with only *Cd274* (+4.3-fold) and *Abca1* (+3.4-fold) significantly upregulated and no genes reaching the threshold for significant downregulation, though *Gas6* (−2.8-fold) and *Pparg* (−2.4-fold) showed notable suppression approaching significance. XDR-MCY infection significantly upregulated *Cd274* (+6.1-fold), *Tyro3* (+5.3-fold), and *Abca1* (+2.9-fold), while significantly suppressing *Cd36* (−3.1-fold), *Gas6* (−3.2-fold1), *Trem2* (−2.8-fold), and *Pparg* (−2.5-fold), the most comprehensive efferocytosis receptor suppression signature across the three strains.

Across all three infection conditions, infection did not broadly enhance the efferocytosis program. Instead, a consistent pattern of selective suppression of apoptotic cell recognition (*Cd36*, *Gas6*) and lipid-sensing (*Pparg*) was observed, alongside apparent upregulation of cholesterol efflux capacity (*Abca1*) and immune checkpoint molecules (*Cd274*). This pattern is consistent with infection-driven rewiring of the macrophage towards a lipid-accumulating, immunosuppressive phenotype that impairs clearance of apoptotic bodies and favors prolonged intracellular bacterial survival.

#### ATRA Activates a Coherent Efferocytosis Program Independently of Infection

Having established that *M. tb* infection suppresses canonical efferocytosis recognition while leaving cholesterol efflux and immune checkpoint pathways paradoxically elevated, we asked whether ATRA activates an efferocytosis program through a mechanism independent of infection-driven signals. This question is important because ATRA-driven efferocytosis induction would need to operate against an infection-conditioned transcriptional landscape to be therapeutically relevant.

In uninfected macrophages, ATRA significantly upregulated *Arg1* (L2FC= +3.30, 9.8-fold), *Abca1* (L2FC= +1.80, 3.5-fold), and *Fasn* (L2FC= +1.69, 3.2-fold), along with directional upregulation of *Stab2* (+7.7-fold), *Abcg1* (+2.5-fold), *Tyro3* (+2.2-fold), *Elmo2* (+1.6-fold), and *Tgfb1* (+1.8-fold). This program was observed entirely in the absence of bacterial challenge, thereby validating that ATRA directly activates efferocytosis gene expression through its own RAR/RXRA transcriptional axis rather than as an indirect consequence of infection-induced signals.

Across infected conditions, ATRA consistently induced *Arg1*, the most robust ATRA-responsive efferocytosis gene in this study in H37Rv+ATRA (L2FC= +1.96, 3.9-fold, padj < 0.001), MDR+ATRA (L2FC= +4.13, 17.5-fold, padj < 0.001), and XDR+ATRA (L2FC= +1.92, 3.8-fold, directional but non-significant). Furthermore, *Abca1* was upregulated in MDR+ATRA (L2FC= +1.08, padj < 0.001), while XDR+ATRA showed no significant efferocytosis gene changes, consistent with the broad transcriptional refractoriness of XDR-MCY-infected macrophages to ATRA identified at the pathway level. The progressive attenuation of ATRA’s efferocytosis-restoring capacity, most robust in uninfected macrophages, partial in H37Rv- and MDR-infected macrophages, and absent in XDR-infected macrophages, parallels the escalating suppression of the translational and metabolic machinery by increasingly drug-resistant strains, suggesting that the severity of infection-driven transcriptional disruption sets a threshold below which ATRA can no longer drive efferocytosis gene induction.

#### RT-PCR and Flow Cytometry Confirm Dose-Dependent ATRA-Driven Efferocytosis Induction at the Protein and mRNA Level

To validate the RNA-seq findings and establish dose-dependence, RT-PCR and flow cytometry was carried out in peritoneal macrophages treated with ATRA at 10 μM and 20 μM for 24 hours (Fig. 5).

**Figure 5.**
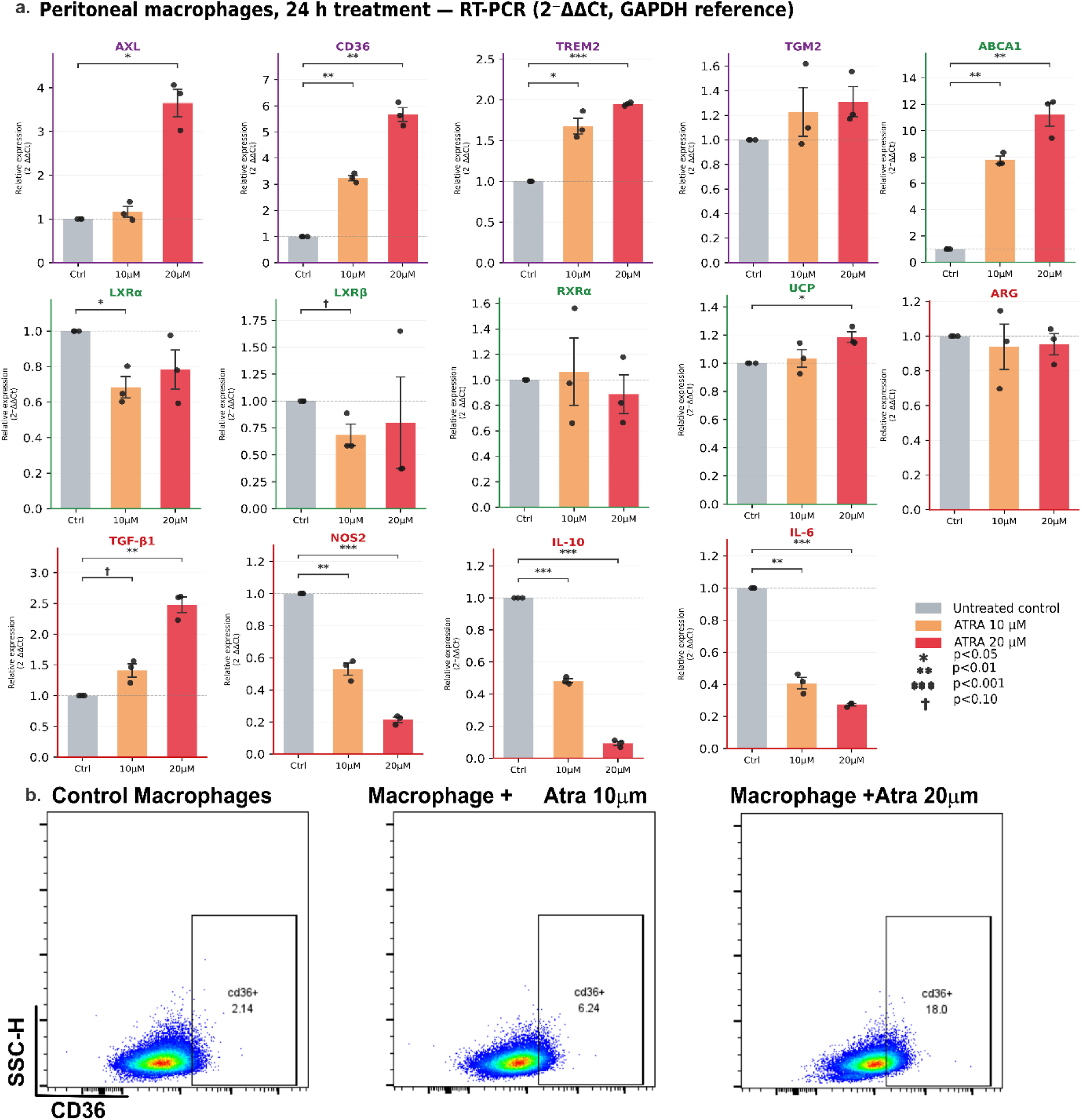
ATRA induces a comprehensive efferocytosis program in uninfected murine macrophages. (a) RT-PCR quantification of efferocytosis-associated gene expression in peritoneal macrophages from C57BL/6 mice treated with ATRA at 10 μM or 20 μM for 24 hours, relative to untreated controls. Expression was calculated by the 2⁻ΔΔCt method using *Gapdh* as the reference gene. Bars represent mean ± SEM (n = 3 independent biological replicates, shown as individual data points). Brackets indicate pairwise comparisons vs untreated control by paired t-test: *p < 0.05, **p < 0.01, ***p < 0.001, †p < 0.10. Dashed horizontal line at y = 1 indicates control expression level. (b) Representative flow cytometry plots showing surface CD36 expression in peritoneal macrophages under untreated conditions (left), ATRA 10 μM (centre), and ATRA 20 μM (right) treatment for 24 hours. Numbers within gates indicate the percentage of CD36-positive cells. Data is representative of three independent experiments.

Flow cytometry demonstrated dose-dependent induction of surface CD36 expression: 2.14% CD36-positive cells in untreated controls, increasing to 6.24% at 10 μM (2.9-fold) and 18.0% at 20 μM (8.4-fold, Fig. 5b). This progressive dose-response at the protein level is consistent with ATRA’s capacity to transcriptionally upregulate *Cd36* as confirmed by RT-PCR and RNA-seq, and thus validates that transcriptional changes translate to biologically meaningful surface receptor remodeling.

RT-PCR validation of 14 efferocytosis and macrophage polarization related genes across the functional panel confirmed robust and dose-dependent effects of ATRA (Fig. 5a). Among efferocytosis receptor genes, *CD36* showed the most pronounced dose-dependent induction (10 μM: 3.24-fold, p = 0.002; 20 μM: 5.68-fold, p = 0.003), while *AXL* showed selective induction at a higher dose (10 μM: 1.16-fold, ns; 20 μM: 3.65-fold, p = 0.014), consistent with the RNA-seq finding of TAM receptor upregulation. *TREM2* was significantly induced at both the doses (10 μM: 1.68-fold, p = 0.019; 20 μM: 1.95-fold, p < 0.001). Within the lipid-handling module, *ABCA1* showed the most striking induction of any gene in the panel (10 μM: 7.78-fold, p = 0.002; 20 μM: 10.88-fold, p = 0.008), reinforcing the centrality of lipid efflux capacity in the ATRA efferocytosis program.

Nuclear receptor and metabolic genes showed more variable responses. *LXRα* was modestly but significantly suppressed at 10 μM (0.68-fold, p = 0.035) with partial recovery at 20 μM (0.78-fold, ns), while *LXRβ* showed a trend towards suppression at 10 μM (p = 0.089). *TGM2*, which encodes transglutaminase 2 involved in apoptotic cell bridging, showed directional but non-significant induction at both doses. *RXRα* and *UCP* did not show significant changes consistent with direct ATRA-driven induction at these doses.

Among the inflammatory and resolution markers, ATRA produced the most statistically compelling dose-dependent effects in the panel. *NOS2*/iNOS was significantly suppressed at both doses (10 μM: 0.53-fold, p = 0.006; 20 μM: 0.21-fold, p < 0.001), *IL-10* was suppressed with striking dose-dependence (10 μM: 0.48-fold, p < 0.001; 20 μM: 0.09-fold, p < 0.001), and *IL-6* was consistently downregulated (10 μM: 0.41-fold, p = 0.004; 20 μM: 0.27-fold, p < 0.001). Concomitant with this anti-inflammatory suppression, *TGF-β1* was significantly induced at 20 μM (2.48-fold, p = 0.007) with a trend towards induction at 10 μM (1.41-fold, p = 0.060), supporting the expected shift from M1 pro-inflammatory to M2 pro-resolution programming. *ARG* showed directional but non-significant changes at the protein quantification level.

Altogether, this data demonstrates that *M. tb* infection actively suppresses canonical efferocytosis receptors and lipid-sensing pathways while paradoxically inducing cholesterol efflux and immune checkpoint genes.

#### *M.tb* Infection Dismantles Efferocytosis Recognition While ATRA Partially Restores it in a Strain-Dependent Manner

Having confirmed that ATRA activates the efferocytosis program in isolation, the study next examined whether *M.tb* infection suppresses this program and whether ATRA can restore it within an infection-conditioned transcriptional landscape. To answer this, and to place efferocytosis within its full immunological context, four functional gene modules were profiled: efferocytosis, M1/IFN signaling, cell death, and immunometabolism/retinoid signaling, across all seven experimental conditions by DESeq2 RNA-seq analysis (Fig. 6).

**Figure 6.**
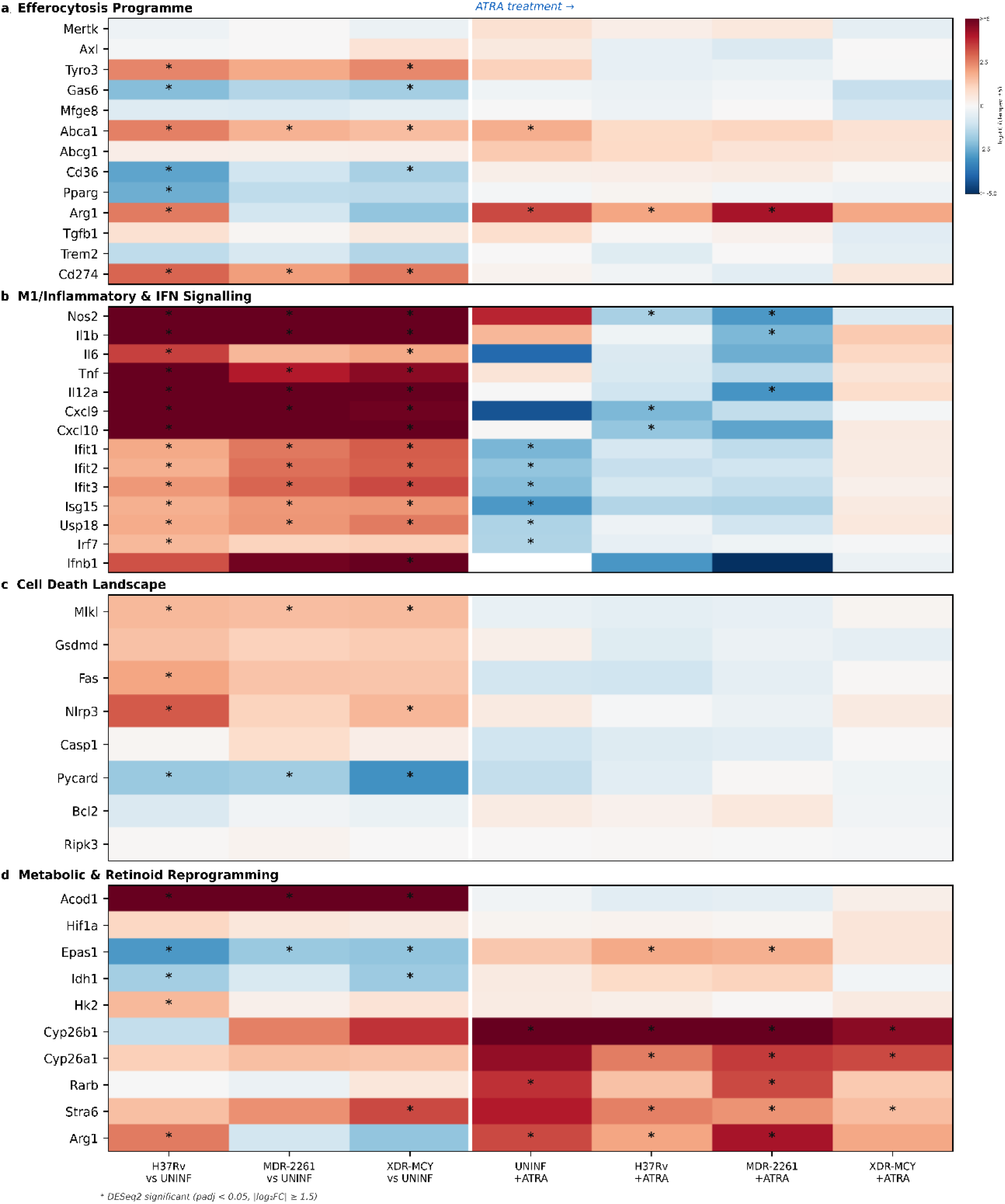
Transcriptional landscape of *M. tb* infection and ATRA recalibration across functional gene modules. Four-panel heatmap displaying L2FC change of key genes across seven pairwise DESeq2 comparisons in peritoneal macrophages: H37Rv vs uninfected, MDR-2261 vs uninfected, XDR-MCY vs uninfected (left three columns, infection response), and UNINF+ATRA vs UNINF, H37Rv+ATRA vs H37Rv, MDR-2261+ATRA vs MDR-2261, XDR-MCY+ATRA vs XDR-MCY (right four columns, ATRA treatment effect). The white vertical line separates infection and ATRA condition groups. Asterisks (*) denote genes meeting DESeq2 significance thresholds (padj < 0.05, |log₂FC| ≥ 1.5). **(a)** Efferocytosis programme: 13 genes spanning TAM receptors (*Mertk*, *Axl*, *Tyro3*), bridging molecules (*Gas6*, *Mfge8*), lipid-handling (*Abca1*, *Abcg1*, *Cd36*, *Pparg*), resolution/M2 mediators (*Arg1*, *Tgfb1*, *Trem2*), and immune checkpoint (*Cd274*). **(b)** M1 proinflammatory and IFN signalling module: cytokines and chemokines (*Nos2*, *Il1b*, *Il6*, *Tnf*, *Il12a*, *Cxcl9*, *Cxcl10*) and interferon-stimulated genes (*Ifit1*, *Ifit2*, *Ifit3*, *Isg15*, *Usp18*, *Irf7*, *Ifnb1*). **(c)** Cell death landscape: necroptosis effectors (*Mlkl*, *Ripk3*), pyroptosis effector (*Gsdmd*), apoptosis mediators (*Fas*, *Bcl2*), and inflammasome components (*Nlrp3*, *Casp1*, *Pycard*).**(d)** Metabolic and retinoid reprogramming: immunometabolic genes (*Acod1*, *Hif1a*, *Epas1*, *Idh1*, *Hk2*) and the retinoic acid signalling module (*Cyp26b1*, *Cyp26a1*, *Rarb*, *Stra6*, *Arg1*). n = 3 per group (UNINF+ATRA n=2).

All three strains universally activated the M1 proinflammatory program (93-100% of module genes significant), with potent induction of *Nos2* (H37Rv: L2FC= +14.0; MDR: +8.7; XDR: +9.8), *Il1b*, *Tnf*, *Il12a*, and *Cxcl9/Cxcl10* (Fig. 6b). IFN signaling was equally universal (86–100% significant), with *Ifit1–3*, *Isg15*, *Usp18*, and *Irf7* significantly induced across all strains. XDR-MCY uniquely drove the highest IFN module mean L2FC(+3.3), anchored by significant *Ifnb1*/IFN-β induction (L2FC= +5.62, padj < 0.001) absent in H37Rv and MDR-2261, a finding of clinical significance given type I IFN’s established role as a tuberculosis susceptibility factor. The metabolic module was robustly activated, with *Acod1* emerging as the most consistently and strongly induced gene across the study (H37Rv: 188-fold; MDR: 75-fold; XDR: 108-fold; all padj < 0.001). Parallelly, *Epas1*/HIF-2α was uniformly suppressed across all strains (H37Rv: L2FC= −2.86; MDR: −1.86; XDR: −1.98; all padj < 0.001) with concomitant downregulation of *Idh1* co-suppressed, indicating that infection drives a HIF-1α–dominant glycolytic shift at the expense of the HIF-2α/Idh1 oxidative resolution axis (Fig. 6d).

Against this backdrop of intense inflammatory activation, infection selectively dismantled efferocytosis recognition (Fig. 6a). The scavenger receptor *Cd36* was significantly suppressed in H37Rv (L2FC= −2.62, −6.2-fold, padj < 0.001) and XDR-MCY (−1.62, padj < 0.001), and the TAM receptor ligand *Gas6* was suppressed across all three strains (H37Rv: −4.4-fold; MDR: −2.8-fold; XDR: −3.2-fold; all padj < 0.001). *Pparg*, coordinating lipid-sensing and efferocytotic transcription, was significantly downregulated across all infections. Paradoxically, *Abca1* and *Cd274*/PD-L1 were induced across all strains, reflecting infection-driven lipid burden and immune checkpoint upregulation, but without the receptor machinery to recognize and clear apoptotic debris. The cell death landscape reinforced this efferocytosis deficit: *Mlkl* was significantly upregulated across all three infection conditions, with *Gsdmd* and *Fas* showing consistent directional but sub-threshold induction (Fig. 6C), collectively indicating a shift toward lytic cell death modes that release bacteria without lysosomal packaging.

In infected macrophages, ATRA consistently and robustly induced *Arg1* across all infection backgrounds (H37Rv+ATRA: 3.9-fold, padj < 0.001; MDR+ATRA: 17.5-fold, padj < 0.001; XDR+ATRA: 3.8-fold, directional), with *Abca1* also significantly upregulated in MDR+ATRA (padj < 0.001). Concurrently, ATRA significantly suppressed *Nos2* in H37Rv-and MDR-infected macrophages, with directional but non-significant suppression of IFN-stimulated genes (*Ifit1*, *Ifit2*, *Ifit3*, *Isg15*) in H37Rv+ATRA; *Ifnb1* was not significantly altered by ATRA in any infected condition (Fig. 6b). ATRA partially reversed the lytic cell death shift, *Gsdmd* suppressed in H37Rv+ATRA and XDR+ATRA, and selectively restored *Epas1* without altering *Hif1a*, defining a novel immunometabolic recalibration axis operating in parallel to efferocytosis (Fig. 6d). The retinoid signaling module confirmed on-target RAR pharmacology across all ATRA conditions through significant induction of *Cyp26b1*, *Cyp26a1*, *Rarb*, and *Stra6*. The net M2−M1 balance shift quantifying ATRA’s overall recalibration was +1.2 for H37Rv+ATRA, +0.9 for MDR+ATRA, and +0.5 for XDR+ATRA (Supplementary Fig. 2), declining with drug resistance and paralleling the gradient of ATRA-responsive pathway enrichment identified in the IPA analysis (23 pathways in uninfected, 3 in MDR+ATRA, 0 in XDR+ATRA).

Taken together, these data establish a coherent mechanistic narrative: *M. tb* infection drives near-complete M1 and IFN activation, shifts macrophage death toward lytic modes that release viable bacteria, suppresses the *Epas1*/Idh1 resolution metabolic axis, and specifically disrupts efferocytosis recognition genes (*Cd36*, *Gas6*, *Pparg*) that would otherwise direct apoptotic bacteria-containing debris toward lysosomal destruction. ATRA counteracts all four dimensions in parallel by restoring efferocytosis receptors, suppressing M1/IFN programming, reversing lytic cell death, and restoring the resolution metabolic axis through on-target RAR/RXRA signaling as validated at the protein and mRNA level in a dose-dependence (Fig. 5). The progressive attenuation of this recalibration with drug resistance identifies the transcriptional permissiveness of the infected macrophage as the limiting factor, and positions MDR-2261 as the context retaining greatest therapeutic potential for ATRA. Given that efferocytosis of *M. tb*-containing apoptotic bodies delivers mycobacteria to the lysosome for bactericidal processing (21), these findings provide a mechanistic rationale for evaluating ATRA as an immunological adjunct in drug-susceptible and MDR tuberculosis.

## Discussion

Using transcriptomic profiling, this study provides a comparative characterization of primary murine peritoneal macrophage responses to drug-susceptible, MDR, and XDR *M.tb* infection and of ATRA-mediated immunomodulation across these clinically distinct resistance profiles. The study reports three important findings. First, infection with all three strains activated a conserved pro-inflammatory program anchored by cytokine storm signaling, cGAS-STING pathway, and potent *Nos2* induction, accompanied by near-complete suppression of host translational machinery, with the magnitude of both activation and suppression escalating progressively with drug resistance. Second, XDR-MCY infection additionally suppressed the antioxidant genes *Prdx1* and *Gpx4*; notably, *Gpx4* suppression was also observed in MDR-2261-infected macrophages, suggesting that antioxidant defense impairment emerges with drug resistance and is most pronounced in XDR infection. Third, ATRA recalibrate infected macrophages toward resolution through induction of efferocytosis program, selective restoration of *Epas1*/*HIF-2α* without disrupting HIF-1α-driven antimicrobial function, and partial suppression of the type I IFN axis, with this recalibration declining progressively with drug resistance.

The conserved macrophage response shared across all three strains was anchored by canonical *M1* activation, suggesting an early host response against *M.tb*. *NOS2*, the major induced genes was robustly upregulated across H37Rv, MDR-2261, and XDR-MCY infection, driving nitric oxide production with established direct antimicrobial activity against *Mtb* through damage to bacterial DNA, proteins, and lipid membranes (22–24). Coordinated induction of *Il1b*, *Il12a*, *Il12b*, *Tnf*, and *Il6* extended the inflammatory program across multiple cytokine axes; notably, *Il1b* drives inflammasome-mediated pyroptosis and immune cell recruitment (25), while *Tnf* is required for granuloma integrity but, when dysregulated, contributes to the tissue pathology characteristic of advanced disease (26, 27). The consistent activation of cGAS-STING signaling (z-score = 2.67-3.89 across strains) reflects cytosolic DNA sensing during active intracellular infection and drives type I interferon induction, a pathway with paradoxical effects in TB, amplifying early innate responses while promoting long-term susceptibility (28). However, upregulation of *Cd274*/PD-L1 across all infection conditions, consistent with prior observations in active TB patients (29), suggests that *M.tb* exploits immune checkpoint signaling to dampen T cell-mediated clearance irrespective of drug resistance profile.

Complementing this inflammatory activation, all three strains imposed near-complete suppression of the host translational machinery including eukaryotic translation elongation and termination, SRP-dependent co-translational protein targeting, NMD, and EIF2 signaling, with z-scores escalating progressively from H37Rv to XDR-MCY. This pattern is consistent with recent evidence that *Mtb* infection triggers stress granule-mediated sequestration of mTORC1, suppressing cap-dependent translation as a mechanism of immune evasion (30). This translational shutdown likely reflects both host metabolic reprioritization toward immune effector programs and bacterial interference with ribosomal function. Beyond translational suppression, all three strains inhibited mitochondrial electron transport chain pathways including oxidative phosphorylation, respiratory electron transport, and Complex I/III/IV biogenesis, with suppression deepening substantially and progressively in MDR-2261 and XDR-MCY compared to H37Rv. This escalating pattern is in concordance with the recent finding, suggesting macrophages with high oxidative phosphorylation (OXPHOS) and low glycolysis harbor reductive drug-tolerant *M.tb* (31). The GAIT translation signaling pathway, which enables selective post-transcriptional silencing of IFN-γ-induced transcripts (32) and is associated with resolution of chronic inflammation, was activated across all three strains but with progressively greater magnitude in MDR-2261 and XDR-MCY, suggesting that counter-regulatory translational pressure on cytokine signaling intensifies with drug resistance, though whether this reflects bacterial exploitation or an adaptive host mechanism to limit immunopathology remains to be determined as tuberculosis is a disease of both infection and inflammation.

The XDR-MCY strain elicited a qualitatively distinct transcriptional signature beyond the quantitative escalation of conserved programs. Most notably, *Ifnb1*/IFN-β induction reached statistical significance exclusively in XDR-MCY, though directional trends were observed across all strains. Concurrently, XDR-MCY was the only strain to significantly suppress *Prdx1*, while *Gpx4* suppression was shared with MDR-2261 but was most pronounced in XDR-MCY. The clinical significance of the *Ifnb1* finding is substantial, as type I IFN signatures correlate with TB progression and susceptibility in human cohorts and murine models (33, 34), with IFN-β specifically driving immunosuppressive IL-10 production and impairing phagosomal maturation (33, 35). XDR-MCY selectively drives this susceptibility axis, representing a potentially important mechanism by which extensively drug-resistant strains circumvent macrophage control. ECM remodeling pathways were activated across all three strains with substantially greater magnitude in XDR-MCY, consistent with the established role of *M.tb*-driven macrophage ECM remodeling in TB immunopathology (36). Notably, the predominant signal in XDR-MCY was activation of collagen biosynthesis and ECM organization rather than degradation, suggesting a fibrotic rather than proteolytic remodeling response that may reflect a distinct macrophage adaptation to extensively drug-resistant infection.

*Acod1* was consistently induced across all three infection conditions, establishing itaconate biosynthesis as a core immunometabolic feature of macrophage-*M.tb* interaction irrespective of drug resistance profile. Itaconate exerts pleiotropic immunomodulatory effects, inhibiting succinate dehydrogenase, restraining inflammatory signaling, and directly limiting bacterial growth (37, 38). However, recent evidence reveals that *ACOD1* plays a dual and opposing role in *M.tb*-infected macrophages. While its enzymatic product itaconate is cytoprotective, high-level ACOD1 expression acts non-catalytically to promote *HSP70* degradation and lysosomal membrane permeabilization, ultimately contributing to macrophage death (39). The parallel induction of *Mlkl* across all infection conditions is therefore consistent with this *ACOD1*-driven lytic death mechanism alongside direct bacterial subversion of the itaconate axis. Notably, ATRA modestly suppressed *Acod1* in H37Rv-infected macrophages but not in MDR or XDR, suggesting that its anti-inflammatory effects operate largely in parallel rather than through the itaconate axis. Alongside *Acod1*, infection-driven suppression of *Epas1*/HIF-2α across all three strains, while *Hif1a* remained unchanged, indicates that *M.tb* infection enforces a HIF-1α-dominant glycolytic program at the expense of the HIF-2α/*Idh1* oxidative resolution axis, an imbalance directly relevant to mechanism of action of ATRA.

Across all infection states, ATRA consistently activated *Cyp26b1* and *Cyp26a1* as canonical RAR targets, with *Stra6* induced in all infected+ATRA conditions and *Rarb* significant in UNINF_RA and MDR_RA contexts, confirming on-target RAR/RXRA engagement (40, 41). Beyond this core retinoid signature, ATRA induced *Abca1* and *Arg1* as the most robustly significant efferocytosis-associated genes (an innate immune effector function) in uninfected macrophages, with directional upregulation of additional efferocytosis components, collectively supporting ATRA-driven engagement of the efferocytosis program through its RAR/RXRA axis (42, 43). This is therapeutically relevant because apoptosis itself is not intrinsically bactericidal and requires subsequent phagocytic uptake and lysosomal fusion of the apoptotic body harboring the bacterium. Efferocytosis delivers *M.tb*-containing apoptotic bodies to the lysosome for bactericidal processing, representing the terminal bactericidal checkpoint following macrophage apoptosis, making it highly effector mechanism of *M.tb*. clearance (21, 44). The consistent induction of *Arg1* across all infected conditions, including the most transcriptionally restrictive XDR-MCY condition, suggests that ATRA retains the capacity to activate efferocytosis-associated signaling even against active drug-resistant infection. Notably, *Arg1* induction was most pronounced in MDR-2261-infected macrophages, substantially exceeding that in H37Rv-infected cells, which may reflect a permissive transcriptional niche created by the intermediate resistance phenotype.

Mechanistically, ATRA showed the selective restoration of *Epas1*/HIF-2α in H37Rv- and MDR-2261-infected macrophages without altering *Hif1a*, that remained unchanged across all infection and ATRA conditions. In macrophages, HIF-1α and HIF-2α are functionally antagonistic rather than redundant. HIF-1α drives the pro-inflammatory Warburg shift, sustains *Nos2*/iNOS-dependent nitric oxide production, and promotes itaconate biosynthesis as part of a broken TCA cycle characteristic of M1 activation (45, 46), while HIF-2α conversely promotes *Arg1*-driven alternative activation and resolution metabolism (47). Infection thus enforces a HIF-1α-dominant pro-inflammatory state by selectively suppressing *Epas1*, and the restoration of *Epas1* by ATRA without altering *Hif1a* re-engages resolution signaling while leaving antimicrobial capacity intact. Notably, efferocytic macrophages stabilise HIF-1α under normoxic conditions through interaction with phosphorylated STAT3 (48), suggesting that ATRA-driven efferocytosis may itself contribute to maintaining *Hif1a* activity, providing a mechanistic rationale for why *Hif1a* remains transcriptionally stable across ATRA conditions despite broad immunomodulatory reprogramming. The functional coupling of HIF-2α and IDH1 in directing carbon flux toward reductive lipid metabolism (49) provides a mechanistic basis for the *Epas1*/*Idh1* co-restoration observed in H37Rv+ATRA, though this co-restoration was attenuated in MDR+ATRA and absent in XDR+ATRA, reflecting the progressive transcriptional refractoriness with drug resistance. Importantly, *Acod1* expression and itaconate biosynthesis were largely preserved under ATRA treatment across all infected conditions. Beyond its antimicrobial role, itaconate alkylates KEAP1 to activate Nrf2, restraining IL-1β, HIF-1α, and type I interferon responses as a negative feedback mechanism to suppress excessive inflammation (50). At high expression levels ACOD1 acts non-catalytically to promote lysosomal membrane permeabilization (39). The preservation of Acod1 under ATRA treatment indicates that its anti-inflammatory effects operate through selective HIF-2α restoration rather than broad metabolic suppression, allowing immunopathology to be tempered without dismantling the core antimicrobial program. Of note, ATRA significantly suppressed *Nos2* in H37Rv- and MDR-infected macrophages but not in XDR-MCY, given the established antagonism between HIF-1α/*Nos2* and HIF-2α/*Arg1* axes (47), these results are consistent with partial M1-to-resolution recalibration where ATRA retains transcriptional permissiveness.

ATRA modestly suppressed interferon-stimulated genes including *Ifit1*, *Ifit2*, *Ifit3*, and *Isg15* in H37Rv-infected macrophages, consistent with its known capacity to attenuate type I IFN signaling (35). As mentioned above, Type I IFN signatures correlate with progressive TB and susceptibility (33), this attenuation is of potential therapeutic relevance, though it was not sustained in MDR- or XDR-infected conditions and *Ifnb1* itself was not significantly altered by ATRA in any infected context. Whether combining ATRA with strategies targeting the type I IFN axis, such as JAK inhibitors, could offer additive benefit in drug-resistant TB warrants further investigation.

## Conclusions

Three principal findings emerge from this macrophage transcriptomic study. First, all three *M.tb* strains drive a conserved inflammatory program anchored by cytokine storm signaling, cGAS-STING activation, and *Nos2* induction, with progressive suppression of host translational and mitochondrial machinery correlating with drug resistance. Second, XDR-MCY imposes a qualitatively distinct immune evasion program characterized by *Ifnb1*/IFN-β induction, suppression of *Prdx1* and *Gpx4*, the latter shared with MDR-2261 and selective downregulation of MHC-II processing genes. Third, ATRA consistently induces *Arg1*-driven resolution signaling across infection backgrounds and selectively restores *Epas1*/HIF-2α in H37Rv- and MDR-infected macrophages without compromising the HIF-1α-driven antimicrobial response or itaconate biosynthesis, a recalibration substantially attenuated in XDR infection. Together, these findings provide a transcriptional rationale for evaluating ATRA as an adjunct to second-line DR-TB regimens, particularly in MDR disease. Future work should prioritize on evaluation of ATRA combined with type I IFN pathway inhibitors for XDR-TB.

## Materials and Method

### Ethics and Animal Care

All animal experiments were conducted in accordance with the guidelines and regulations of Department of Biotechnology (India) and were approved by the Institutional Animal Ethics Committee of International Centre for Genetic Engineering & Biotechnology (ICGEB), New Delhi. Female C57BL/6 mice (6-8 weeks old) were used for ex- vivo studies conducted at Tuberculosis Aerosol Challenge Facility of ICGEB under standard animal care and handling protocols.

### Bacterial Strains

*M.tb* strain H37Rv (American Type Culture Collection 25618) was obtained from the Colorado State University repository. The MDR-Jal2261 (resistant to ethambutol, isoniazid and rifampicin) and XDR-MYC-431 *M.tb* strains were a generous gift from Dr. KVS Rao (ICGEB). All mycobacterial strains were grown in 7H9 (Middlebrooks, Difco^TM^, USA) medium, broth supplemented with 10% oleic acid, albumin, dextrose and catalase (OADC) at 37°C and maintained in the Biosafety Level 3 facility in ICGEB. The cultures were maintained till the mid-log phase and were then aliquoted in 20% glycerol and stored at −80 °C. These cryopreserved stocks were subsequently used for all experiments.

### Peritoneal Macrophages: Isolation, Reagents & Treatment

C57BL/6 mice were intraperitoneally injected with 4% sodium thioglycolate to induce peritonitis. Peritoneal macrophages were isolated on day 5, seeded in 6 well plate and incubated overnight. On day 6, the macrophages were infected with different *M.tb* strains (H37RV, MDR-2261, XDR-MCY) for 4 hours with MOI (Multiplicity of infection) 1:10. After four hours of infection, cells were washed with 1x sterilized PBS followed by addition of RPMI-1640 media supplemented with 10% fetal bovine serum (51, 52). Cells were then treated with retinoic acid (10μM) for 24 hours. Cells without Retinoic acid treatments were taken as a control. The experimental design included three technical replicates: Uninfected Macrophages (Control); Uninfected Macrophages treated with ATRA; Macrophages infected with *M.tb* H37RV; infected with *M.tb* H37RV and treated with ATRA; infected with MDR strain of *M.tb*; infected with MDR strain of *M.tb* and treated with ATRA; infected with XDR strain of *M.tb*; and infected with XDR strain of *M.tb* and treated with ATRA.

### RNA Extraction, Library Preparation and Sequencing

After 24 hours of treatment, cells from control, experimental and ATRA-treated groups were collected, washed twice with PBS, and pelleted by centrifugation. Total RNA was extracted using the RNeasy Mini Kit (QIAGEN, Cat. No. 74104) and quantified using a NanoDrop spectrophotometer (Thermo Scientific). Poly-T oligo-attached magnetic beads were used to capture the poly-A tails of mRNA molecules from the total RNA sample. The purified mRNA was then fragmented, and cDNA was synthesized in two steps: first-strand synthesis using random hexamer primers, followed by second-strand synthesis. Final library was prepared through a series of steps including end repair, A-tailing, adapter ligation, size selection, amplification, and purification. Sequencing was conducted by Novogene using the Illumina NovaSeq PE150 platform based on paired-end sequencing technology to generate 150-bp reads.

### Quality Filtering, Alignment, and Quantification

Quality assessment of the raw transcriptomic data was performed using FastQC (v0.11.9) (https://www.bioinformatics.babraham.ac.uk/projects/fastqc/). The reads were then trimmed using Trimmomatic 0.39 (53), applying the quality filters (LEADING:3, TRAILING:3, SLIDINGWINDOW:4:15, MINLEN:36). The cleaned reads were then aligned to the GRCm39 mouse reference genome (GCF_000001635.27) using STAR aligner (v2.7.10b) (54). The STAR alignment was performed using parameters optimized for spliced alignments, with filters for multi-mapping reads and strict criteria for identifying intron-spanning reads. Finally, gene expression quantification was performed using RSEM (v1.3.3) (55), which generated gene-level count data for subsequent analyses.

### Differential Expression and Pathway Analysis

Differential expression analysis was performed using DESeq2 v1.38.3 (56) in R v4.3.1. To address the potential confounding effect of bacterial strain variation (H37Rv, MDR-2261, XDR-MCY), two complementary design strategies were implemented. For pairwise comparisons between infected and uninfected samples within each strain (e.g., RV vs UNINF, MDR vs UNINF, XDR vs UNINF), a simplified design formula (∼ group) was employed, as strain is inherently confounded with infection status in these comparisons. Similarly, for direct strain-strain comparisons (MDR vs RV, XDR vs RV, XDR vs MDR) and for within-strain ATRA treatment effects (UNINF_RA vs UNINF, RVATRA vs RV, MDR_RA vs MDR, XDR_RA vs XDR), ∼ group design was used.

To isolate the global ATRA treatment effect across all infected samples while accounting for strain-specific expression differences, a multi-factor design with strain as a covariate: ∼ strain + ATRA was used. This model pools all infected samples (RV, MDR, XDR with and without ATRA treatment, n=18) and estimates the ATRA effect adjusted for baseline differences among the three *M.tb* strains. Statistical significance was determined using the Wald test with Benjamini-Hochberg (BH) false discovery rate (FDR) correction. Genes with *Padj* < 0.05 and |L2FC| ≥ 1.5 were considered differentially expressed.

The identified DEGs were then analyzed using Ingenuity Pathway Analysis software (IPA; QIAGEN) to perform pathway analysis and construct Protein-Protein Interaction (PPI) networks. Pathways or networks were considered significant if they met two criteria: an absolute Z-score ≥ 2 and an adjusted p-value (padj) < 0.05.

### Multi-Pathway Transcriptional Analysis and Efferocytosis Gene Profiling

Four functional gene modules were defined *a priori* based on established pathway annotations: efferocytosis programme (23 genes) (42–44), M1 proinflammatory and IFN signalling (14 genes) (43, 57, 58), cell death (8 genes) (59), and immunometabolic/retinoid reprogramming (10 genes) (40, 60) (Supplementary Table 8). Differential expression across all seven pairwise comparisons was performed using DESeq2 (v1.38.3) with BH FDR correction; significance thresholds were similar as stated previously. L2FC values for all module genes were assembled into a genes and conditions matrix, with values clamped at ±5 for display to prevent high-magnitude outliers from compressing the colour scale for biologically informative mid-range changes. The fig.s were generated in Python (matplotlib). To quantify overall M1→M2 recalibrating effect of ATRA, a net M2 − M1 balance score was computed for each ATRA condition as the difference between mean L2FC of the M2/efferocytosis module and the mean L2FC of the M1 proinflammatory module.

For surface protein validation, macrophages treated with ATRA (10 μM and 20 μM, 24 h) were stained with fluorochrome-conjugated anti-mouse CD36 and anti-mouse CD274/PD-L1 antibodies (BioLegend) followed by analysis through flow cytometry (Cytek Aurora). Real-time quantitative (RT-PCR) analysis was performed using Bio-Rad Real-Time thermal cycler (Bio-Rad, USA). The reaction was set up according to the manufacturer’s instruction. The enzyme reverse transcriptase and SYBR green Master mix were obtained from Bio-Rad. The relative expression level of mRNAs was normalized to that of internal control GAPDH by using 2-ΔΔCt cycle threshold method. The primers used for the study (Supplementary Table 9) were purchased from Integrated DNA Technologies (IDT), USA. For statistical comparisons, unpaired Welch t-test was used

## DATA AVAILABILITY

RNA sequencing data generated in this study are deposited in the NCBI Sequence Read Archive under BioProject ID PRJNA1201576.

## ACKNOWLEDGEMENTS

This research was partially supported by the Science and Engineering Research Board (SERB) under sanction order PDF/2021/001238. MK thanks DHR-ICMR for the funding (R.11013/49/2021-GIA/HR). RK gratefully acknowledges financial support from the University Grants Commission Ba-sic Scientific Research Start-Up Grant (No. F.30-527/2020(BSR)).

---

**Supplementary Figure 1.**
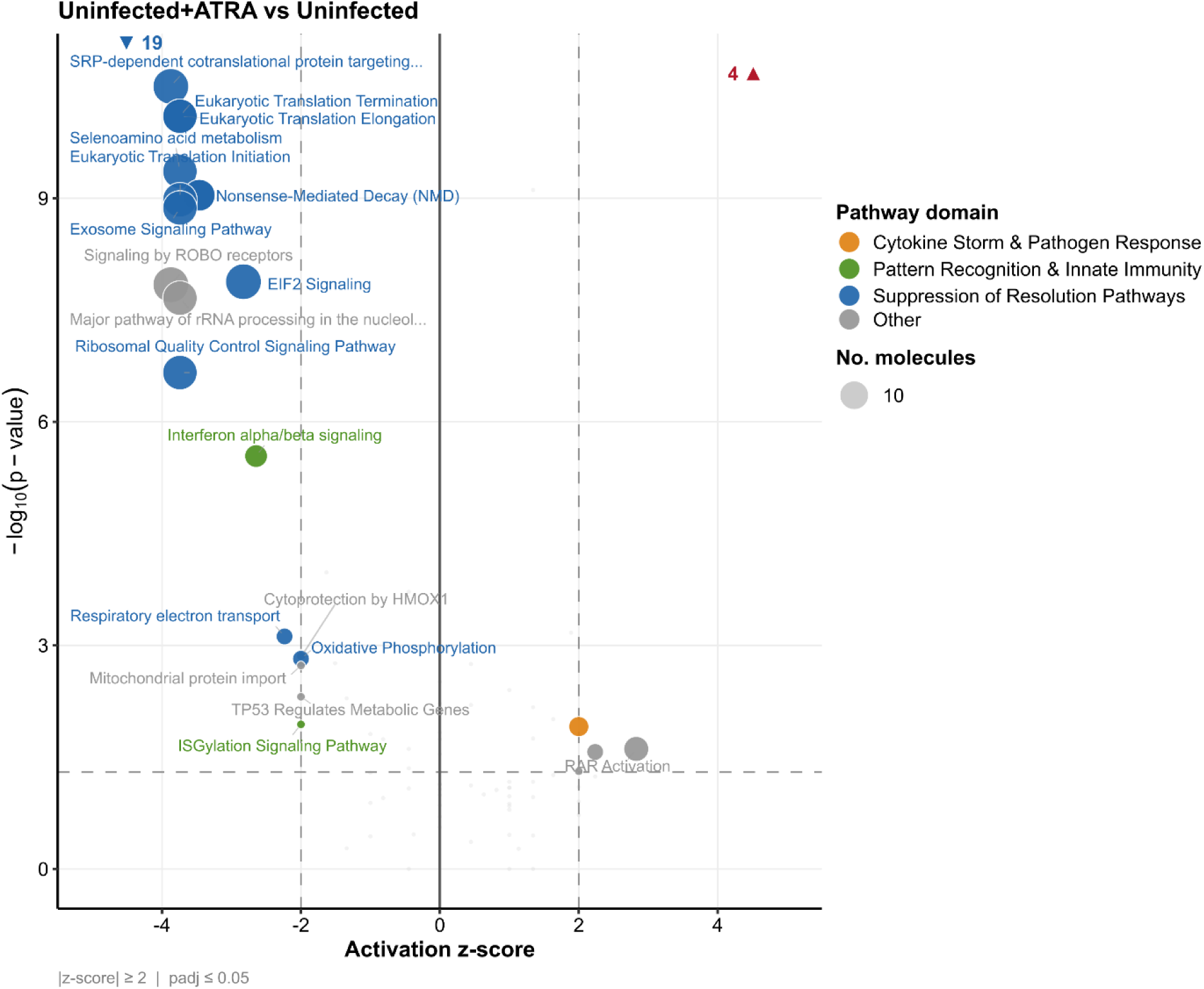
IPA canonical pathway analysis of ATRA treatment in uninfected macrophages. Scatter plot depicting significantly dysregulated canonical pathways in uninfected macrophages treated with all-trans retinoic acid (ATRA) compared to untreated uninfected controls (n = 2 vs 3; note: UNIFATRA2 excluded following QC failure due to abnormally low library depth). Each dot represents one IPA canonical pathway. The x-axis shows the activation z-score, where values ≥ 2 indicate pathway activation and values ≤ −2 indicate inhibition relative to the untreated condition. The y-axis represents the −log₁₀(p-value) of pathway enrichment. Dot size is proportional to the number of dataset molecules mapping to the pathway. Dot colour denotes the biological domain: orange, cytokine storm and pathogen response; green, pattern recognition and innate immunity; blue, suppression of resolution pathways; grey, other. Non-significant pathways (|z-score| < 2 or p > 0.05) are shown as light grey background points.

**Supplementary Figure 2.**
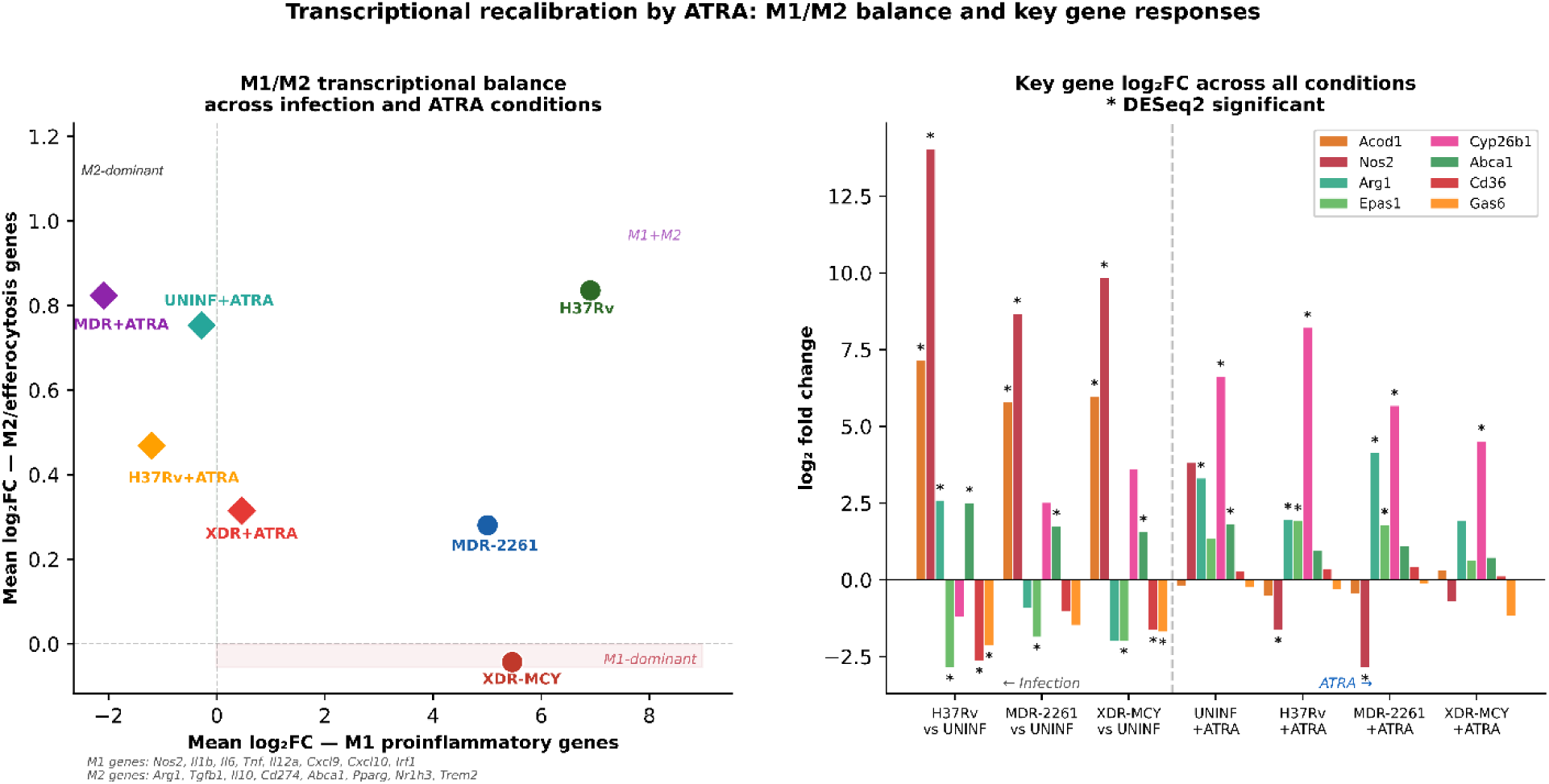
Transcriptional recalibration by ATRA: M1/M2 polarization balance and key gene responses across infection states. **(Left) M1/M2 transcriptional balance scatter.** Each point represents one experimental condition plotted by its mean L2FC of M1 proinflammatory genes (x-axis) versus mean L2FC of M2/efferocytosis genes (y-axis), calculated across all genes within each module with a valid L2FC estimate. Circles denote infection conditions (H37Rv, MDR-2261, XDR-MCY vs uninfected); diamonds denote ATRA-treated conditions (UNINF+ATRA, H37Rv+ATRA, MDR+ATRA, XDR+ATRA vs respective untreated controls). Dashed lines at x = 0 and y = 0 indicate no net change relative to the reference condition. M1 module genes: *Nos2*, *Il1b*, *Il6*, *Tnf*, *Il12a*, *Cxcl9*, *Cxcl10*, *Irf1*. M2/efferocytosis module genes: *Arg1*, *Tgfb1*, *Il10*, *Cd274*, *Abca1*, *Pparg*, *Nr1h3*, *Trem2*. **(Right) Key gene L2FC bar chart.** L2FC of eight representative genes, spanning immunometabolism (*Acod1*, *Epas1*), M1/M2 polarisation (*Nos2*, *Arg1*), efferocytosis (*Abca1*, *Cd36*, *Gas6*), and retinoid signalling (*Cyp26b1*), across all seven pairwise DESeq2 comparisons. The dashed vertical line separates infection comparisons (left, three conditions) from ATRA treatment comparisons (right, four conditions). Asterisks (*) denote genes meeting DESeq2 significance thresholds (padj < 0.05, |log₂FC| ≥ 1.5, BH-FDR correction). n = 3 per group; UNINF+ATRA n = 2.

